# Biogeographic multi-species occupancy models for large-scale survey data

**DOI:** 10.1101/2021.11.05.467527

**Authors:** Jacob B. Socolar, Simon C. Mills, Torbjørn Haugaasen, James J. Gilroy, David P. Edwards

## Abstract

1. Ecologists often seek to infer patterns of species occurrence or community structure from survey data. Hierarchical models, including multi-species occupancy models (MSOMs), can improve inference by pooling information across multiple species via random effects. Originally developed for local-scale survey data, MSOMs are increasingly applied to larger spatial scales that transcend major abiotic gradients and dispersal barriers. At biogeographic scales, the benefits of partial pooling in MSOMs trade off against the difficulty of incorporating sufficiently complex spatial effects to account for biogeographic variation in occupancy across multiple species simultaneously.
2. We show how this challenge can be overcome by incorporating pre-existing range information into MSOMs, yielding a ‘*biogeographic multi-species occupancy model*’ (bMSOM). We illustrate the bMSOM using two published datasets: Parulid warblers in the United States Breeding Bird Survey, and entire avian communities in forests and pastures of Colombia’s West Andes.
3. Compared to traditional MSOMs, the bMSOM provides dramatically better predictive performance at lower computational cost. The bMSOM avoids severe spatial biases in predictions of the traditional MSOM and provides principled species-specific inference even for never-observed species.
4. Incorporating pre-existing range data enables principled partial pooling of information across species in large-scale MSOMs. Our biogeographic framework for multi-species modeling should be broadly applicable in hierarchical models that predict species occurrences, whether or not false-absences are modeled in an occupancy framework.

## Introduction

Community ecologists often seek inference about occurrence patterns of multiple species simultaneously. To improve inference, especially for infrequently detected species, models such as the multi-species occupancy model (MSOM) share information across species via hierarchical random effects (Devajaran et al 2020). The MSOM was originally developed for application to relatively homogenous study areas where occupancy probabilities vary little across space (Dorazio & Royle 2005). Subsequently, the MSOM has been applied to a variety of landscapes where occupancy probabilities are modeled as a function of site-specific covariates (e.g. Tingley & Beissinger 2013, Rich et al 2017, Ribeiro Jr. et al 2018).

At very large spatial scales that subsume biogeographic variation in species ranges, the biologically relevant covariate structure becomes exceedingly complex. The biogeographic distributions of multiple species depend on species-specific interactions between numerous environmental, geographic, and historical factors. Rather than attempting to parameterize and fit these myriad effects, large-scale single-species distribution models often eschew the generalized linear modeling framework in favor of highly flexible additive effects (Rushing et al 2020) or machine-learning methods such as MaxEnt (Phillips et al 2006) or regression trees (Fink et al 2010). Such approaches, however, are not easily amenable to pooling information across data-poor species for robust community-level inference. As a result, there is a need for hierarchical multi-species approaches to study community variation across biogeographic spatial scales (Janousek & Drietz 2020; Jarzyna & Jetz 2018).

When the data at hand are insufficient to estimate realistic parametric models that adequately capture the biogeography of every species in a study region, the benefit of partial pooling across species trades off against the detriment of species-specific spatial biases. Across complex biogeographic landscapes, models that fail to robustly account for species’ ranges will tend to underestimate occupancy at points within a species’ range and overestimate occupancy at points outside a species’ range.

Given the difficulty of estimating sufficiently flexible spatial effects in large-scale MSOMs, previous authors have applied a variety of *post hoc* strategies to address inferential problems that arise from fitting simple MSOMs to biogeographically complex regions. For example, Jarzyna & Jetz (2018) applied a MSOM to predict terrestrial bird richness across the coterminous United States and manually adjusted their model output by setting occupancy probabilities to zero in regions where a species does not occur. Janousek & Drietz (2020) applied a MSOM to the spatial complex bird communities of the greater Rocky Mountains of the United States, but 30 species (29%) failed a posterior predictive check and were excluded from further analysis. However, such statistical palliatives are not sufficient to ensure reliable inference, because *post hoc* exclusion of species or geographic ranges still allows poorly modelled data to inform inference in the remainder of the model.

Never-observed species pose additional modeling challenges at biogeographic scales. In principle, traditional MSOMs can handle never-observed species via data augmentation with excess pseudospecies, each of which is given an all-zero detection history and is included or excluded from the true community according to a Bernoulli random variable with modeled probability \omega (Dorazio & Royle 2005). However, these models cannot estimate independent covariate relationships for the never-detected species, and some authors have chosen to exclude data-augmented pseudospecies from downstream analyses (e.g. Tingley & Beissinger 2013). Incorporating data-augmentation approaches into models that leverage traits or phylogeny to predict detection (Sólymos et al 2017) or occupancy (e.g. via trait-environment interactions) is especially challenging, requiring potentially dubious assumptions about the trait distributions for never-detected species or the discretization of traits into functional guilds (Tenan et al 2017).

Recent progress towards multi-species pooling in biogeographic-scale MSOMs, including models for never-observed species, has focused on discretizing the study region into spatial units (Tobler et al 2015, Sutherland et al 2016) and discretizing the community into ecological guilds (Tenan et al 2017). Sutherland et al (2016) propose a multi-region model where data-augmented MSOMs are fit to each region and region-specific community richness is directly modeled as a function of covariates. Importantly, the identities of species, including species that are never detected in a particular region, are fixed across regions, thus enabling pooled estimation of species-specific occupancy and detection probabilities across the entire multi-region study area. Tobler et al (2015) fit a similar model without data augmentation, such that the potential pool of species in any region is exactly the total pool of species observed study-wide, and all species identities are fixed and known. Tenan et al (2017) extend the approach of Sutherland et al (2016) to trait-based models, discretizing the community into ecological guilds and estimating the richness of never-observed species for each guild separately. However, all of these methods rely on the assumption that, conditional on covariates, the spatially discrete regions are internally homogeneous and mutually independent. Therefore, they are not suitable for application to biogeographic landscapes with complex and continuous spatial variation.

For many taxa, a wealth of pre-existing biogeographic information is available in the form of range maps, geospatial sightings databases, and/or published range descriptions. We hypothesized that by leveraging this information, we could develop simple, tractable multi-species models that yield reliable pooled inference about in-range occupancy probabilities while avoiding the pitfall of conflating in-range and out-of-range occupancy probabilities within and across species. We achieve such inference by collapsing complex, multidimensional biogeographic variation into simple summary covariates, which we call *range covariates*. Possible range covariates include (transformations of) the distance to the nearest geographic range margin or elevational range limit. Such information is increasingly available, especially for taxa amenable to occupancy modeling (Jetz et al 2012). We refer to an MSOM that incorporates range covariates as a *biogeographic multi-species occupancy model* (bMSOM). Like the multi-region model of Tobler et al (2015), the bMSOM fixes the identity of every species (including never-observed species) in the dataset. However, unlike all previous models, the bMSOM handles arbitrarily complex biogeographic-scale spatial dependencies using a very simple covariate structure.

Here, we formally describe the bMSOM and apply it to two published datasets: 51 Parulid warbler species in the United States Breeding Bird Survey (49 observed; 2 never-observed), and 910 bird species in forests and pastures of Colombia’s West Andes (314 observed, 596 never-observed). In addition to providing a mechanism for principled data pooling across very large spatial scales, bMSOMs fix the identity of every species in the metacommunity and link those identities to real-world species with known traits. They are therefore exceptionally suited to trait-based models for occupancy and detection, analyses of point-scale richness, and biogeographically pooled analyses of the influence of local-scale environmental variation on community composition or structure. Furthermore, bMSOMs sometimes allow for *a priori* exclusion of data at extralimital sites (where occupancy is implausible), thereby reducing the total dataset size and the computational resources required for model fitting.

## Methods

### Model formulation

We formulate the likelihood for the standard MSOM as

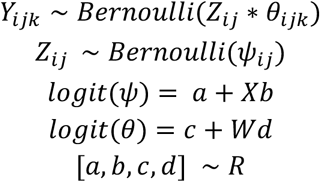

where *i*, *j*, and *k* index the species, site, and visit; *Y*_*ijk*_ is the binary detection/non-detection data; *Z*_*ij*_ is the latent true occupancy state; *θ*_*ijk*_ is the detection probability conditional on occupancy (i.e. *p*(*Y*_*ijk*_ = 1 | *Z*_*ij*_ = 1)); *ψ*_*ij*_ is the occupancy probability (i.e. *p*(*Z*_*ij*_ = 1)); *a* and *c* are column vectors of intercepts for occupancy and detection, respectively; *X* and *W* are design matrices for occupancy and detection, respectively; *b* and *d* are column vectors of coefficients for occupancy and detection, respectively; and *R* is the joint random effects distribution for *a*, *b*, *c*, and *d*. At a minimum, *R* must include random intercepts by species for both occupancy (*a*) and detection (*c*).

For computational efficiency, we re-express the model by marginalizing over the latent occupancy states *Z*, and for models without visit-specific detection covariates we replace the Bernoulli likelihood indexed over visits with a binomial sufficient statistic for the total number of detections at a point.

### The bMSOM

The likelihood for the bMSOM is no different from the standard MSOM; what differs is the data. As in a data-augmented MSOM (Dorazio & Royle 2005), we append all-zero detection histories for never-observed species. However, each of these all-zero entries corresponds to a specific species that we know *a priori* occurs in the biogeographic vicinity of the sampling points, and so we treat all species as present in the metacommunity and “available” for occupancy.

Crucially, we include one or more “range covariates” that describe whether a given species is in-range or out-of-range at each point, and we estimate species-specific random coefficients for the range covariates. For example, if species-specific minimum and maximum elevation data are available along an elevational gradient, an appropriate range covariate might be the squared elevational distance from a survey point to the midpoint of a species’ elevational range. If species differ substantially in their elevational breadth, we might rescale these differences for each species separately prior to squaring, such that values of 1 correspond to the species-specific upper range limits and values of −1 correspond to the species-specific lower range limits. When species are distributed over two-dimensional space rather than along one-dimensional gradients, we suggest using a range covariate based on the geographic distance from a survey point to the species geographic range margin. When only crude range descriptions are available, the range covariate might simply be binary, designed to distinguish areas that are clearly out-of-range. Regardless of the precise nature of the range covariates, we include them in the bMSOM with species-specific random slopes.

In the bMSOM context, it is sometimes additionally useful to completely exclude severely out-of-range species-site combinations from analysis, a process that we call “*biogeographic clipping”*. By excluding sites where occupancy probabilities are *a priori* negligible, it is possible to improve within-range estimation while reducing the computational burden of model fitting. For example, biogeographic clipping can account for sharp range margins associated with abrupt biogeographic barriers (e.g. mountains or deepwater marine barriers) while still allowing occupancy probabilities to decay more gradually at range margins elsewhere. Likewise, migratory species can be modeled with temporal clipping, where species-site combinations are removed from the data if the site was surveyed outside of the dates of potential presence. Here we assume that repeat visits to a site occur sufficiently quickly to avoid problems of closure for migratory species.

### Example 1: warblers of the coterminous United States

Jarzyna & Jetz (2018) analyzed multi-species occupancy patterns of North American birds by applying a traditional MSOM to the United States Breeding Bird Survey (BBS) dataset (Bystrak 1981). With a focus on the year 2018, the coterminous United States, and the Parulid warblers, we reimplemented the modeling framework of Jarzyna & Jetz (2018) and compared the traditional MSOM to the bMSOM. We restricted the analysis to Parulid warblers (as opposed to the full North American avifauna) for the sake of computational efficiency, and we selected the Parulid warblers in particular because they are relatively speciose (51 species breed in the coterminous US), display marked variation in species’ ranges, are well sampled by BBS protocols, and are sufficiently behaviourally homogeneous to ensure that they approximately satisfy exchangeability assumptions for hierarchical modeling, even in the absence of species-specific covariates.

We downloaded BBS data for the year 2018 from www.pwrc.usgs.gov, and we obtained range maps for all Parulid warblers that regularly breed within 200 km of the coterminous United States from Birdlife International (BirdLife International and Handbook to the Birds of the World 2019). To develop a range covariate, we measured the distance from the starting point of each BBS survey route to the nearest edge of each species’ range, excluding range limits associated with shorelines. We then sought a transformation of the species ranges that would approximately linearize the logit-proportion of occupied points. We believed *a priori* that the function should asymptote at large negative distances (i.e. in the core of the range), and we sought a function that would asymptote at zero, such that hierarchical model components would effectively be setting a prior on occupancy probabilities in the core of a species range. We believe that an asymptote at zero should help to ensure exchangeability across species and should aid in eliciting informative priors (if desired).

We binned all BBS point-species combinations by their distance to the range edge (negative distances at in-range points, positive distance at out-of-range points), and we examined several functions to select one that approximately linearizes the logit-proportion of occupied points, ultimately selecting the inverse logit of distance-to-range expressed in units of 200 km (Supporting Information, section 1).

We fit three occupancy models to these data. Model 1 is the model of Jarzyna & Jetz (2018), a traditional MSOM with correlated random intercepts for detection and occupancy and a random slope for the effect of elevation on occupancy (Zipkin et al 2009). Model 2 is a bMSOM that extends Model 1 via the inclusion of the range covariate described above. Model 3 is the bMSOM with biogeographic clipping, equivalent to model 2 but excluding all species-point combinations >400 km from the mapped range, a distance beyond which no detections existed in the data.

We compared the predictive performance of models 1, 2, and 3 via approximate leave-one-out cross-validation using Pareto-smoothed importance sampling with moment-matching (Vehtari et al 2017, 2021). For each pair of models, we compared overall predictive performance as well as predictive performance for each species separately. For models 1 and 2, we compared predictive performance over all points as well as over just the subset of points that we retained after biogeographic clipping. For comparisons involving model 3, we evaluated predictions only over the subset of points retained after biogeographic clipping, as this model sees nondetections outside of the clipped range as deterministic structural zeros.

### Example 2: Forest conversion in Colombia’s West Andes

We applied the bMSOM to a dataset of bird species at 146 point-count stations on an elevational gradient from 1260 to 2680 masl in Colombia’s West Andes. Each point was visited on four consecutive or nearly-consecutive days (Gilroy et al 2014a,b). Points were located in either forest or pasture and were arranged in clusters of three points each nested inside one of three subregions. Following the taxonomy of BirdLife International, 910 bird species potentially occurred in the vicinity of the points, based on biogeographic clipping (see below). Of these, 314 were detected at least once and 596 were never observed. Our inferential goal was to assess how point-scale species richness varies along the elevational gradient.

We implemented a biogeographically clipped bMSOM framework to address Colombia’s exceedingly complex biogeography. We incorporated two range covariates, based on elevation and geography. The elevational range covariate was based on the elevational limits reported in Ayerbe Quiñones (2018), supplemented with several additional references for species whose taxonomic treatment differed between Ayerbe Quiñones (2018) and BirdLife International (2020). We standardized the elevations of each point across species by linearly rescaling the raw elevations of the points such that an elevation of 1 corresponded to the upper range limit, and an elevation of −1 corresponded to the lower range limit (Figure S1). We implemented biogeographic clipping at species-standardized elevations of −3 and 3, beyond which our dataset contained no detections. We additionally implemented temporal clipping for migratory species, treating all migrants as deterministically absent outside of their normal dates of presence in Colombia (Supplementary Data).

We derived the geographic range covariate using digital range maps from Ayerbe Quiñones (2018; see also Vélez et al 2021). We implemented biogeographic clipping at a buffer of 160 km around these maps, as well as at the crest of the West Andes and the floor of the Cauca Valley to address the complex biogeography of birds in this region. Against this biogeographic clipping, our field data exposed errors of omission in just 2 species, indicative of the high quality of the Ayerbe Quiñones maps. For these two species, we added range around previously known clusters of records (eBird 2021) that coincided with the records in our data, and we refined the biogeographic clipping to incorporate an appropriate buffer around this additional range (see Supporting Information, section 2). The spatially balanced sampling of forest and pasture ensures that this *post hoc* addition of range does not compromise inference about responses to deforestation. Our field data exposed no errors of omission against our temporal clipping.

After performing all clipping, we selected an appropriate transformation of raw distance for the geographic range covariate following the procedure described for warblers above. Again, we approximately linearized the logit-proportions by taking the inverse logits of the distance, this time measured in units of approximately 14.9 km (Supporting Information, section 1).

We modeled occupancy on the logit scale based on an intercept and coefficients for the geographic range covariate, the elevational range covariate (linear and quadratic terms), interactions between the elevational range covariate (linear and quadratic terms) and whether the species occurs at lowland elevations, land-use, 19 species traits (Table S1), and the interactions of those 19 traits with land-use. Our random effects structure incorporated random taxonomic intercepts for species and family, random spatial intercepts for species:cluster and species:subregion, and random taxonomic coefficients for all range covariates (species-specific terms) and for land-use (species- and family-specific terms).

We modeled detection on the logit scale based on an intercept and coefficients for land-use, time-of-day (given as hours post-sunrise), four species traits, and the interaction between time-of-day and the median elevation where a species occurs. Our random effects structure incorporated random taxonomic intercepts for species and family, a random intercept for species:observer, and random taxonomic coefficients for time-of-day (species-specific terms) and land-use (species- and family-specific terms). See Supporting Information (section 3) for details of our prior specification.

To assess our ability to recover principled trait-based estimates of sensitivity to deforestation, including for never-observed species, we compared the sensitivity estimates from the model against independently estimated forest dependency scores from BirdLife International (2020). We repeated this comparison for just species with at least one observation in our data and for just the never-observed species.

We then use the bMSOM to predict the local species richness (including never-observed species) in forest and pasture across an elevational gradient in the Colombian West Andes. For comparison, we also estimated species richness along the elevational gradient in forest and pasture using a data-augmented multi-species occupancy model including the 314 observed species and 1000 never-observed pseudospecies (Dorazio & Royle 2005). To enable use of the data-augmented model, we removed all species-specific covariates (including information about range, traits, dates of occurrence, and family-level classification) from the analysis.

### Model fitting

We implemented occupancy models using Hamiltonian Monte Carlo sampling in Stan (Stan Development Team 2021) via R packages brms (Bürkner 2017) for the Parulid warblers and cmdstanr (Gabry & Češnovar 2021) for the Colombian birds. We performed model comparison using the R package ‘loo’ (Vehtari et al 2020). For all warbler models, we ran 4 chains for 1000 warmup iterations and 1000 sampling iterations. For the Colombian Andes, we ran 4 chains for 1500 warmup iterations and 1500 sampling iterations. For the data-augmented model, we encountered substantial challenges in model fitting; we describe these problems and their resolution in the Supporting Information (section 4). We ensured that all models (except the data-augmented model; see Supporting Information, section 4) converged with maximum r-hats less than 1.03 for all parameters, no divergences in the Hamiltonian trajectories, and energy fraction of missing information greater than 0.2 in all chains.

## Results

### Warblers of the coterminous United States

The inclusion of the distance covariate in the bMSOM yielded dramatic improvements in predictive performance (Figure 1), with an improvement in expected log predictive density (ELPD) of 8558 with standard error (SE) 127. Predictive performance improved for 50 out of 51 species, the only exception being a marginal decrease of −0.4 (SE 0.8) for Tropical Parula (Figure 1a).

**Figure 1:**
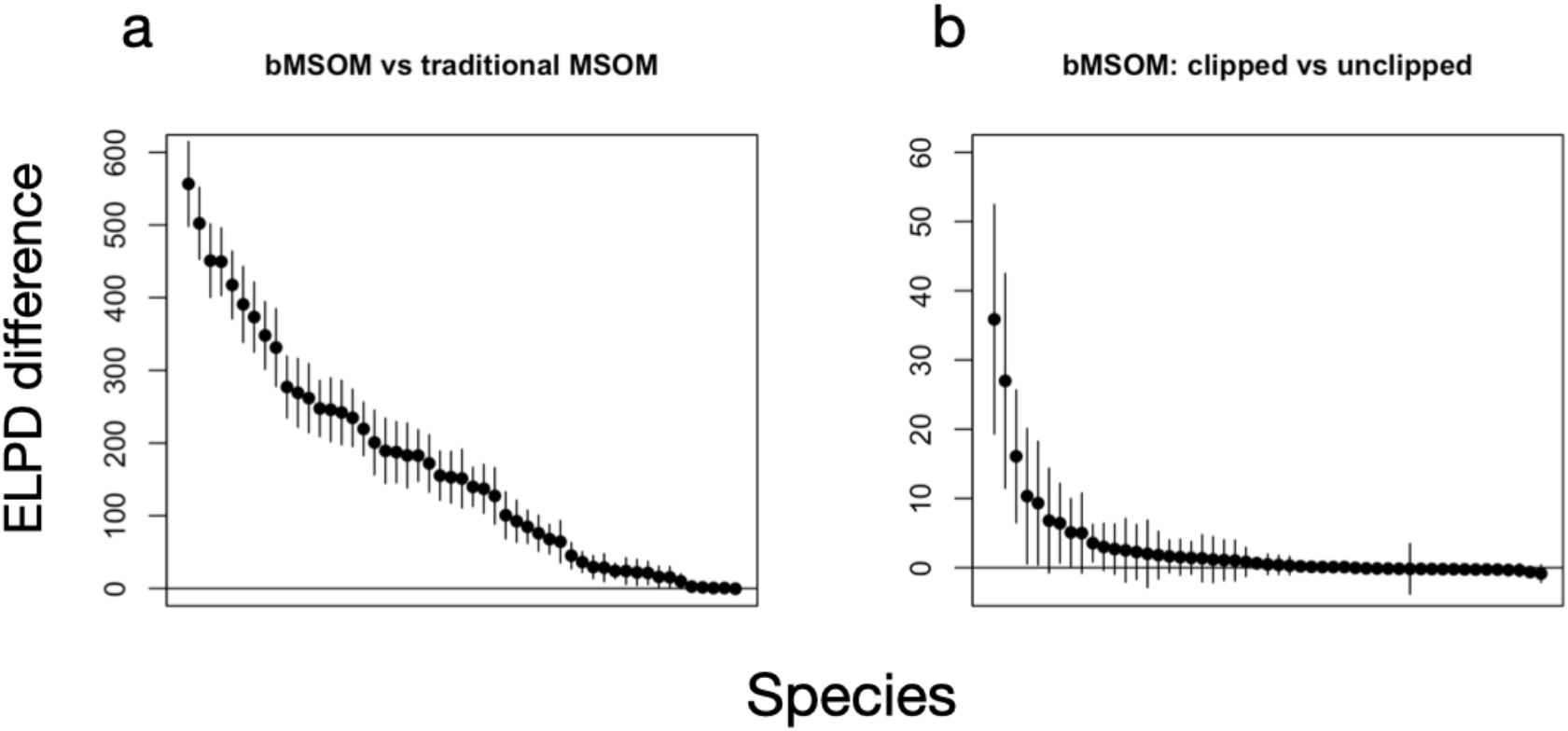
Differences in species-specific expected log pointwise density (ELPD) calculated by approximate leave-one-out cross validation for the BBS data. Points represent species-specific posterior means and are ordered by decreasing ELPD difference; lines represent +/− 2 standard errors. Positive values indicate superior predictive performance in the first model. Comparisons are performed across all points (a) or across just the subset of points that are retained in the clipped model (b).

Biogeographic clipping delivered further gains in predictive performance at in-range points, with an ELPD improvement of 147 (SE 17). Within the species-specific regions that were retained after clipping, predictive performance improved for 33 out of 51 species (Figure 1b). Among the 18 species that did not see improvements, we observed the largest decrease in ELPD in Lucy’s Warbler, but even this decrease was only marginal (−0.9; SE 0.6), which is too small to conclude that the clipped model performs worse than the unclipped model for Lucy’s Warbler or any other species. Biogeographic clipping also yielded substantial gains in computational efficiency, reducing the runtime by a factor of almost three, from a mean per-chain execution time of 5.8 hours (worst-case chain 6.1 hours) to 2.0 hours (worst-case chain 2.1 hours) on an M1 Macbook Air.

Predicting the traditional MSOM across space yielded relatively uniform occupancy probabilities compared to the bMSOM, which universally predicted higher occupancy probabilities at in-range points and lower probabilities at out-of-range points (Figure 2). The predictions of the clipped bMSOM were generally quite similar to those of the unclipped bMSOM, though small differences were apparent for some species. In these cases, the clipped model tended to estimate steeper elevational relationships, which reflects the clipped model’s flexibility to fit locally appropriate relationships unconstrained by the need for accurate prediction at severely out-of-range sites. For example, the Mourning Warbler is restricted to the eastern United States, and within this range it tends to occur at high elevations. Without clipping, the bMSOM estimates only a modest positive elevational relationship (1.4, 95% CI 0.8–2.0), because steeper estimates yield unacceptably high occupancy probabilities in the high-elevation mountains of the western United States (Figure 3). Clipping allows the model to estimate an appropriately steep relationship (3.2, 95% CI 2.1–4.4) within the species’ northeastern range. We provide maps of predicted occupancy probabilities for all species in Figure S2.

**Figure 2:**
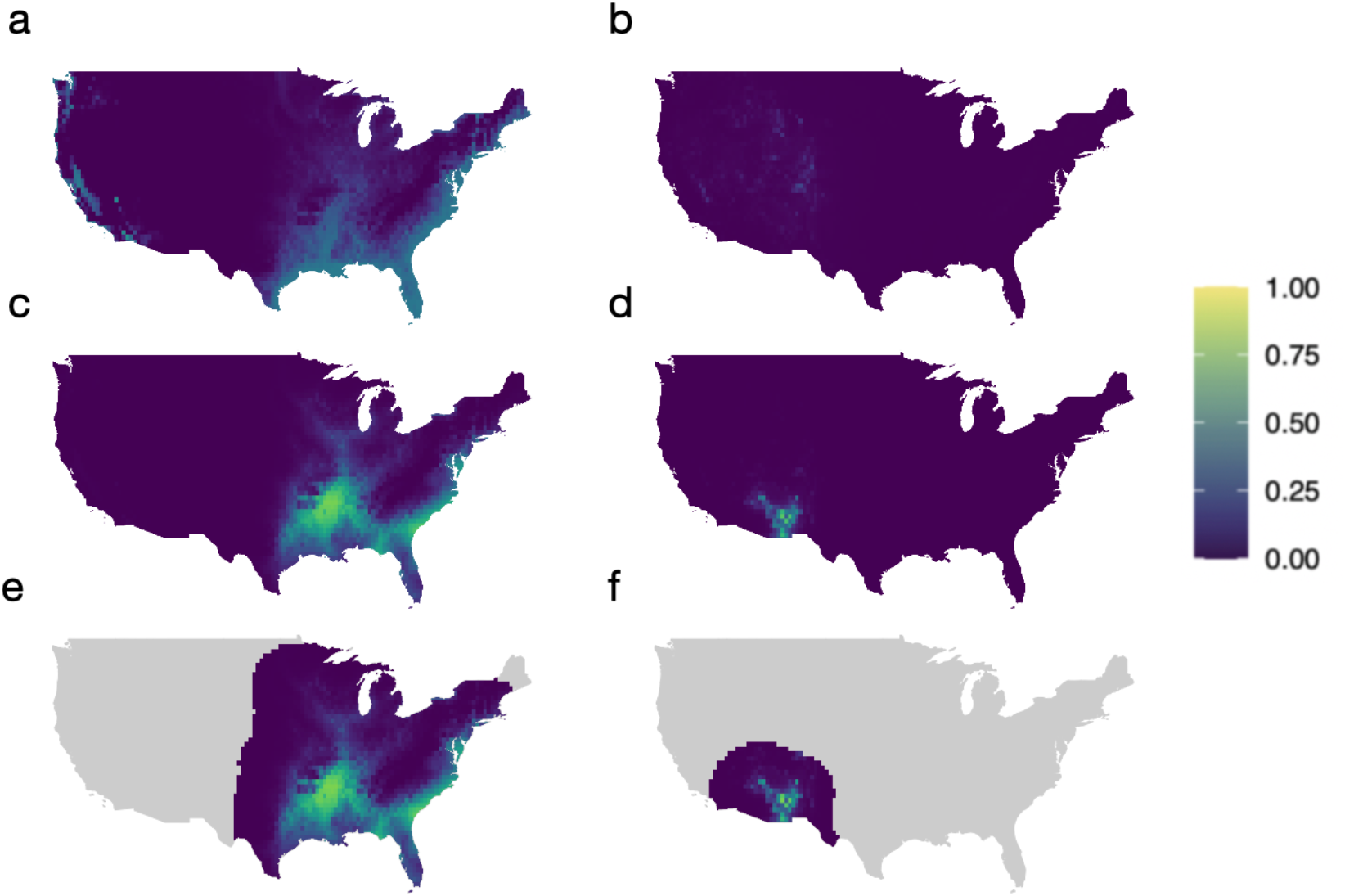
Predicted occupancy probabilities for Prothonotary Warbler (left-hand column), a low-elevation eastern species, and Red-faced Warbler (right-hand column), a high-elevation southwestern species. Predictions are given by the traditional MSOM (a, b), the bMSOM (c, d), and the clipped bMSOM (e, f). Equivalent figures for all species are available in Figure S2.

**Figure 3:**
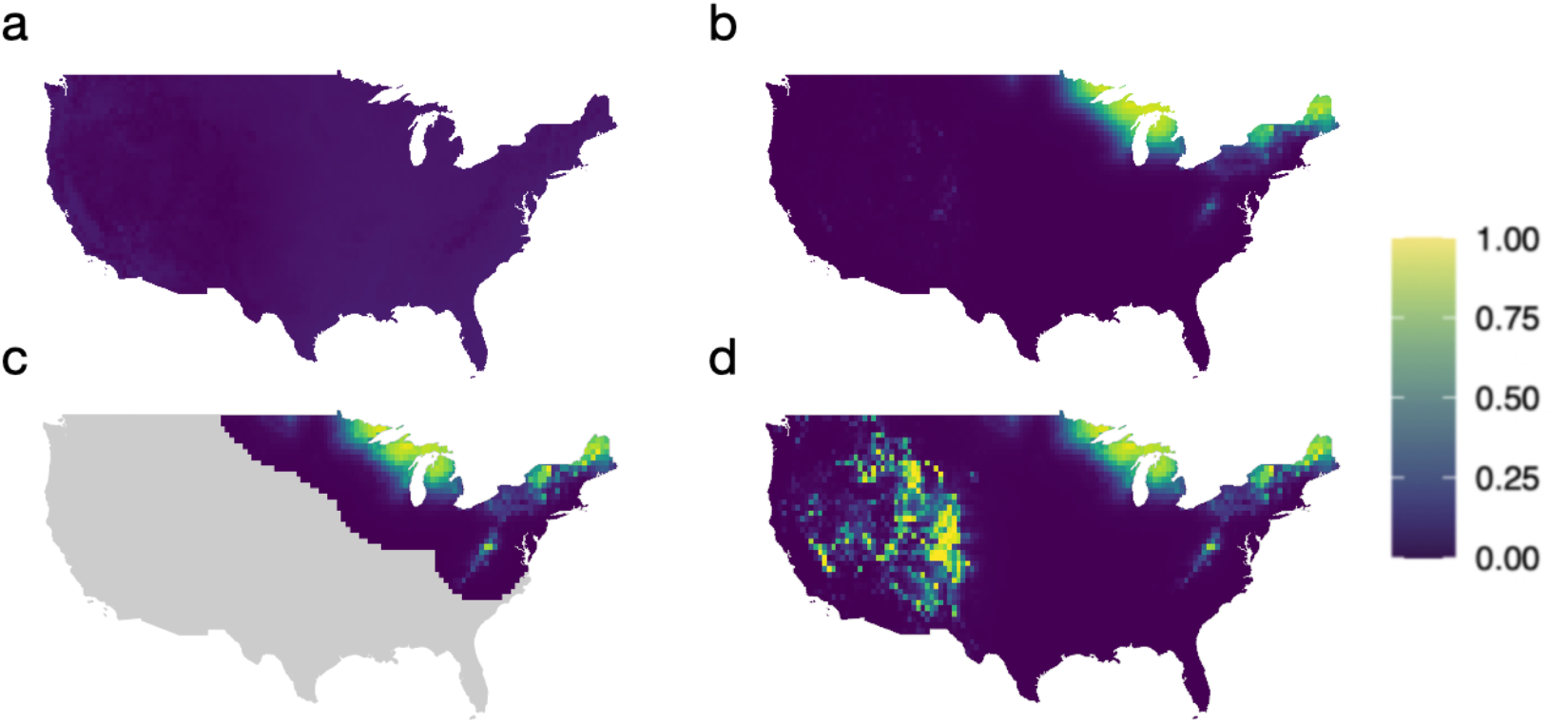
Predicted occupancy probabilities for Mourning Warbler from a) the traditional MSOM, b) the bMSOM without clipping, c) the clipped bMSOM, and d) the clipped bMSOM coefficients projected across the entire country. Mourning Warbler tends to occur at higher elevation within its eastern range. Biogeographic clipping allows the model to estimate appropriately strong elevational relationships in the eastern United States, without the need to avoid predicting high occupancy probabilities across the mountainous western United States. In (d) the steep elevation relationship overcomes the negative relationship with geographic distance to yield high occupancy probabilities in the mountains of the western United States.

### Forest conversion in Colombia’s West Andes

The bMSOM yielded reliable trait-based inference for the sensitivity of the entire avifauna, including never-observed species (Figure 4). We provide a summary of the fitted model posterior in Table S1. Among species with at least one observation and classified by BirdLife International as having either high forest dependence or low/no forest dependence, the model universally inferred that species classified as highly forest-dependent responded more negatively to forest conversion than other species. Even among species with no observations, the model successfully inferred that the vast majority of species classified as having high forest dependence respond more negatively to pasture than species with low/no forest dependence (Figure 4). Moreover, some of this overlap may arise due to the difficulty of accurately categorizing forest dependence in rarely encountered species.

**Figure 4:**
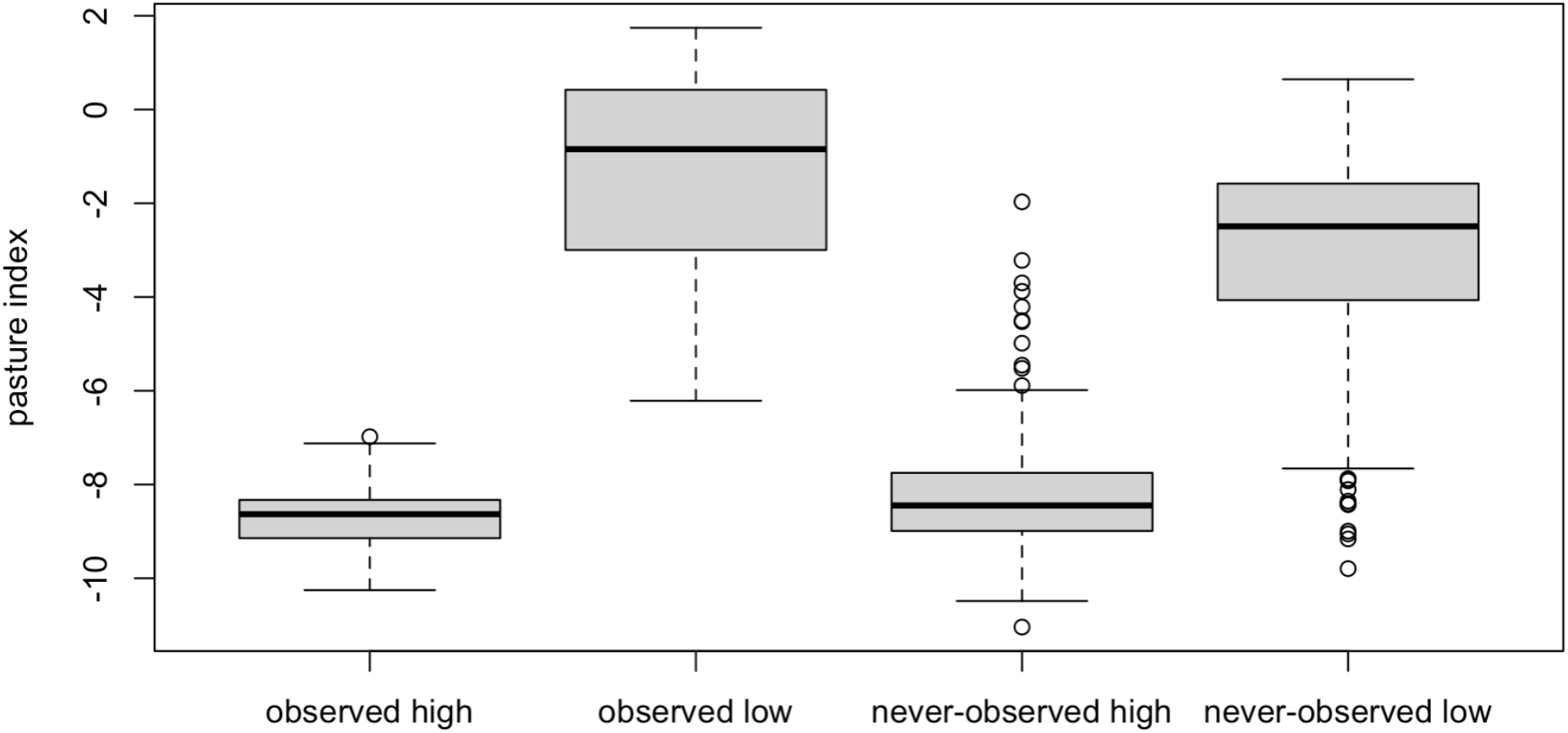
Model-estimated logit-scale effects of forest conversion to pasture on avian occupancy in the West Andes. The pasture index is the difference in logit-scale occupancy probability between forest and pasture. Data points are posterior medians for individual species. Box plots are grouped by whether or not the species was detected at least once during sampling (observed/never-observed), and by whether BirdLife International independently ascribes the species to the “high” forest dependency category (high) or to the “low” or “not a forest species” categories (low). Species ascribed to “medium” or “unknown” forest dependency categories are omitted from this figure.

The bMSOM provided species-specific trait-based inference on occupancy probabilities in both pasture and forest, even for never-observed species. While the data-augmented MSOM predicted similar patterns of alpha-diversity along the gradient (Figure 5a,b), the data-augmented model also displayed specific pathologies that affect both its practicality as an inferential tool and the quality of the resulting inference. First, the data-augmented MSOM required dramatically more computational resources and fine-tuning of algorithmic parameters to successfully fit (Supporting Information, section 4). Second, the data-augmented model implausibly estimated that species were included in the metacommunity with probability near unity (95% credible interval 0.995-1.000). The data-augmented model accounts for the non-detection of the 1000 never-observed pseudospecies species by ascribing to them extreme elevational ranges that overlap little with the sampling points. Thus, the data-augmented model predicts that never-observed species occur most frequently at both the lower and upper extremes of the gradient (Figure 5c,d). At the lower end of the gradient, this pattern is expected based on the tendency for species richness to increase with increasing productivity and forest stature towards lower elevations and is consistent with the predictions of the bMSOM. At the upper end, however, this pattern is at odds with both theoretical expectations and with the predictions of the bMSOM. In particular, the data-augmented model estimates a spurious increase in alpha richness near the highest sampling points (all of which are in forest; Figure 5a) due to an uptick in occupancy of never-observed species (Figure 5c).

**Figure 5:**
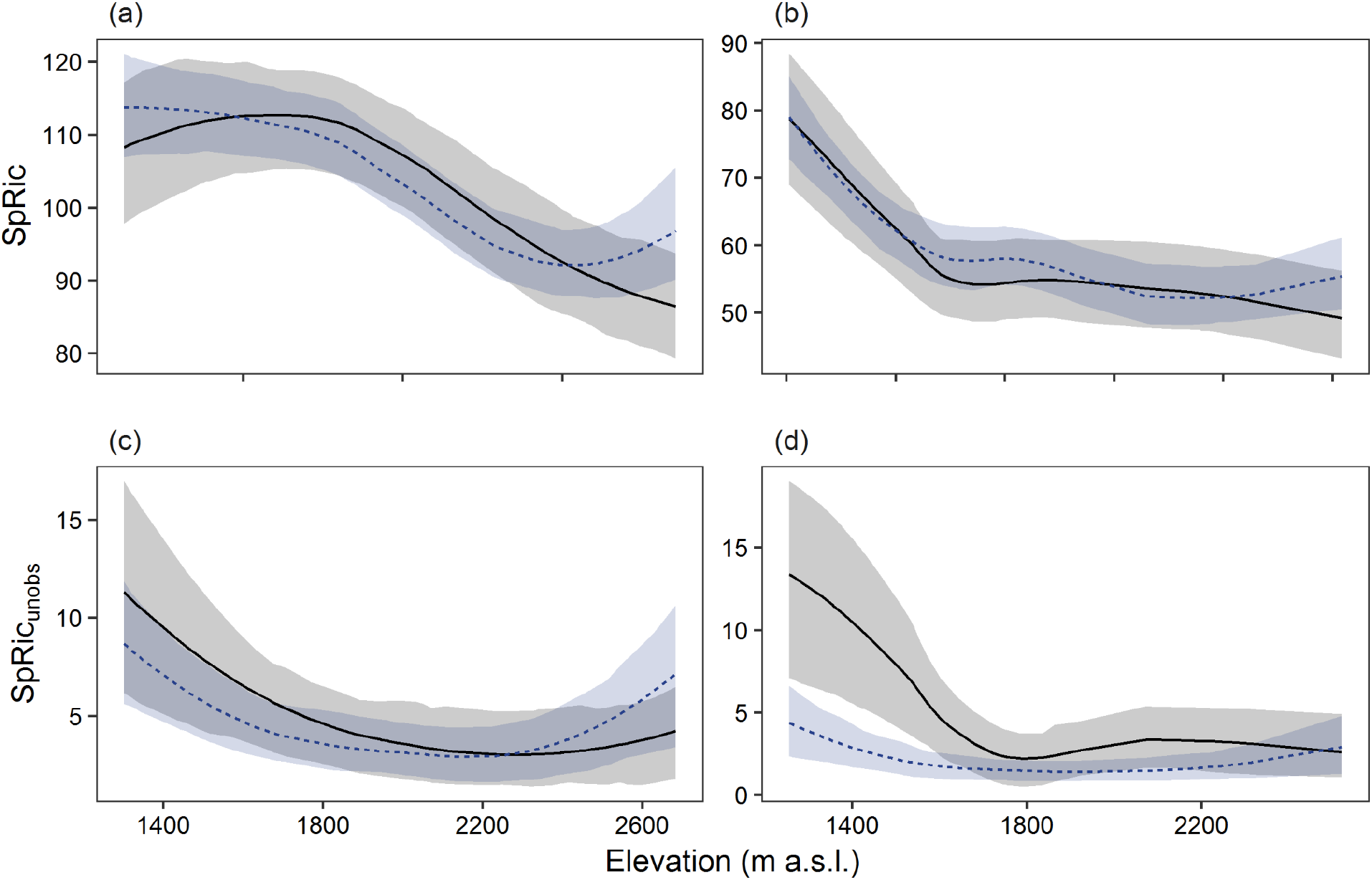
Predicted point-scale avian richness for all species (a,b) and only never-observed species (c,d) along the West Andean elevational gradient in forest (a,c) and pasture (b,d). The bMSOM is shown in gray and the data augmented MSOM is shown in blue with 80% credible interval overlaid. The data-augmented model spuriously infers an excess of never-observed species at points on the upper end of the elevational gradient.

## Discussion

Ecologists increasingly seek inference about occurrence patterns for multiple species over vast spatial scales (Jarzyna Jetz 2018, Janousek & Dreitz 2020). We show that it is possible to leverage the existing multi-species occupancy model framework to provide principled inference over these large scales simply by including covariates that summarize the positions of sampling points with respect to species ranges. When appropriate range information is available, the resulting biogeographic models deliver improved predictive performance, trait-based inference for unobserved species, and computational speed-up compared to traditional approaches. The application of traditional MSOMs (that lack range covariates) at biogeographic spatial scales results in lack-of-fit and unrealistic spatial predictions, suggesting that modelers must proceed with great caution when fitting large-scale MSOMs in contexts where *a priori* range information is unavailable.

### Core advantages of the bMSOM

There are three main advantages of bMSOM over traditional MSOMs. First, we get better inference on the observed species for a range of reasons. It is unsurprising that the introduction of a covariate that reliably distinguishes in-range from out-of-range points should improve predictive performance, but the full scope of the improvement is broad and at times subtle. For example, although revising extralimital occupancy probabilities to zero after model fitting eliminates obviously erroneous predictions of extralimital occupancy, doing so does not enable accurate estimation of within-range occupancy probabilities (Jarzyna & Jetz 2018). The traditional MSOM’s conflation of occupancy probabilities at in-range and out-of-range points induces a strong negative bias in occupancy probabilities at in-range points (Figure 2). Range covariates additionally improve predictive performance by placing species on a common scale, where exchangeability assumptions are more likely to hold. For instance, by using species-standardized elevations to model avian occupancy in Colombia’s West Andes, we ensured that the magnitudes of the quadratic coefficients are likely to be similar across species, irrespective of heterogeneity in elevational range breadth. In traditional MSOMs, heterogeneity in elevational range breadth might be confounded with phylogeny, traits, or other predictors of interest, which could impede clear inference about covariate relationships (Sólymos 2017).

Likewise, the geographic range covariates ensure that the intercepts for all species correspond to occupancy probabilities in their core ranges, partially removing potential relationships between intercepts and range size. In the BBS analysis, the bMSOM yielded large gains in predictive performance even for the most widespread species in the dataset, the Common Yellowthroat (ELPD gain 153, SE 18). Part of this effect results from principled handling of range margins in southern Texas and California, but part arises from better pooling of the intercept across species. Whereas the traditional occupancy model pools the intercept for Common Yellowthroat with intercepts for other species that reflect a mixture of in-range and out-of-range occupancy probabilities, the bMSOM pools the intercept for Common Yellowthroat with intercepts that reflect in-range occupancy probabilities across all species. Thus, the bMSOM estimates a larger intercept for Common Yellowthroat than the naïve MSOM, yielding better predictive performance.

A second key advantage of incorporating range covariates into occupancy models is their ability to specify both the metacommunity size and the identity of every species in the metacommunity. Doing so simplifies the formulation and implementation of models that include never-detected species, enables the use of species-specific covariates to model occupancy for never-detected species, and eliminates uncertainty in the total metacommunity size as a source of uncertainty in point-level species richness. Thus, the bMSOM ameliorates the substantial computational challenges associated with fitting the data-augmented model, the limitations on inference about never-detected species inherent to the data-augmented model (Tingley & Beissinger 2013), and a variety of pathologies that arise in the data-augmented model when detection probabilities are low (see Tingley et al 2020). At the same time, the bMSOM avoids assuming that variation in occupancy across space or in trait distributions affecting occupancy or detection is readily discretized (Sutherland et al 2016, Tenan et al 2017). In our analysis of the West Andean avifauna, the bMSOM was able to recover the forest dependency of never-observed bird species with high fidelity (Figure 4). By leveraging this ability, we were able to predict alpha-richness in forest and pasture for the full community using a procedure that avoided predicting spurious patterns among never-observed species (Figure 5) and was not subject to the computational challenges of fitting the data-augmented model (Supporting Information, section 4).

A third benefit associated with range covariates is the ability to perform biogeographic clipping, which can substantially reduce the computational burden of model fitting (a three-fold reduction in our BBS analysis) and can improve model fit at biologically relevant sites (Figure 3). Biogeographic clipping also enables modelers to account for biogeographic barriers that produce abrupt drops in occupancy probability, with zero occupancy probability on one side of the barrier. Such drops are difficult to capture with general-purpose range covariates that must also account for the more gradual decay in occupancy probability at other range margins. By removing species-point combinations on the wrong side of biogeographic barriers from analysis, these out-of-range detection histories do not propagate (mis)information about the distance-decay in occupancy probabilities near mapped range margins elsewhere.

### Application in practice and conclusions

The importance of prior range information highlights one potential pitfall in occupancy modelling (including the traditional occupancy model) at scale: when is a range map good enough? While a range map does not need to precisely reflect range margins, significant errors of omission will carry through to posterior inference with zero occupancy probabilities at locations where a species is in fact present. In our West Andes dataset, for example, we identified deficiencies for a minority of species (n=2), requiring manual adjustment of range maps to bring them up to date with known species’ occurrences. While expert knowledge can be harnessed both to assess the quality of range maps as well as make any requisite changes, this does raise the danger of inflated “researcher degrees of freedom” (Simmons et al 2011), and manual updating of range maps based on the observed data. Judicious care is needed to ensure that these choices do not generate unfounded inference.

Overall, the bMSOM carries advantages that are especially well suited for estimating covariate relationships by pooling across species with disparate ranges, uncovering local-scale covariate relationships while controlling for broad-scale biogeography, estimating alpha-scale species richness, and trait-based modeling of never-observed species. On the other hand, due to the requirement for pre-existing range data the bMSOM is ill suited for exploratory species-distribution modeling at biogeographic scales or inference about the effects of environmental predictors that are spatially autocorrelated over scales comparable to species entire ranges. We caution, however, that except in data-rich contexts where ranges can be reliably estimated from data for all species under study (and thus multispecies approaches are unlikely to be necessary or useful), traditional approaches that do not incorporate range information are likely to yield poor inferences about occupancy, biasing in-range occupancy probabilities downwards while also predicting non-negligible occupancy in many areas far removed from a species’ range. There thus appear to be general limitations to the application of MSOMs at large spatial scales that subsume significant biogeographic turnover. In the absence of major sampling efforts that allow range-wide variation in occupancy to be estimated from the data directly, application of occupancy models to species or taxonomic groups for which range information cannot be included will typically yield poor inference.

## Acknowledgments

We thank Oswaldo Cortes, who assisted with data collection in Colombia, and volunteers for the US Breeding Bird Survey for producing the datasets analyzed here. We thank the community of Stan users, and particularly Sebastian Weber and Bob Carpenter, for useful pointers regarding the implementation of our models. We thank Fernando Ayerbe Quiñones and the Instituto de Investigación de Recursos Biológicos Alexander von Humboldt for facilitating our access to range maps This work was funded by the Research Council of Norway (project no. 262378) and the Natural Environment Research Council (grant number NE/R017441/1).

## Author contributions

JBS and SCM conceived of the method and analyzed the data. JJG, DPE, and TH provided data for the West Andes. All authors contributed to writing the manuscript and approved of the final version. JBS and SCM contributed equally to this work.

## Data availability

All data and code will be deposited in Figshare upon acceptance of the manuscript. For review purposes, code is available at https://github.com/jsocolar/bmsom_paper.

## Supporting Information

### Section 1: Choosing a transformation for the distance covariate

The relationship between distance-to-range and occupancy probability is not expected to be logit-linear. In particular, we expected *a priori* that the occupancy probability in the core range of wide-ranged species would asymptote at a value less than one. We recognized *a priori* that by selecting a transform that mapped strongly negative (i.e. in-range) distances to the vicinity of zero, we would yield a model where the pooling of species-specific intercepts shared information about the core of the range for all species. Thus we initially considered transformations of the form *e*^*x*^, where *x* is the distance-to-range and the measurement unit for *x* is freely chosen by the researcher. However, these transforms failed to approximately linearize the logit-proportion of points with at least one detection (aggregated across all species).

We then experimented with transforms of the form 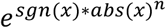, where *sgn* is the sign function, *abs* is the absolute value function, and the exponent *n* is freely chosen by the researcher. This extra degree of freedom allowed us to approximately linearize logit-proportions choosing *n* to be near 0.7, but at the cost of transforming the largest positive (out-of-range) distances to extremely large values. As a consequence, this transform resulted in an over-abundance of extremely influential points during model fitting, which made model comparison via approximate leave-one-out cross-validation using pareto-smoothed importance sampling prohibitive. Moreover, we realized after the fact that this transformation requires the estimation of relationships that are *a priori* implausible since in some years (but not in 2018) the breeding bird survey records a species thousands of kilometers out-of-range.

Thus, we finally settled on an inverse-logit transformation of distance, with distance expressed in units of 200 km for the BBS data and units of 14.92 km in the West Andes data (this unit is not round because when we chose it we were working with the scaled raw distances, i.e. the distances divided by their standard deviation). Again these are freely chosen values to approximately linearize the logit-proportions of points with at least one detection, aggregated across all species.

### Section 2: Manual range-map updates in the West Andes avifauna

Our field data exposed only two errors of omission in the biogeographic clipping that we performed around the range maps of Ayerbe Quiñones (2018):

#### Entomodestes coracinus

We detected this species on the east slope of the West Andes in a region where Ayerbe Quiñones maps this species as being (barely) restricted to the west slope (thereby triggering biogeographic clipping at the crest of the West Andes). Our records coincide with numerous additional east-slope records, and we manually added polygons around known clusters of east-slope records (eBird 2021).

#### Thlypopsis superciliaris

We detected this species at multiple locations. Ayerbe Quiñones omits this species from the northern part of the West Andes in Colombia, despite its well documented occurrence in the area. We manually added a polygon around a well-known cluster of records in the northern West Andes (eBird 2021).

We then recomputed the distance-from-range and reimplemented biogeographic clipping based on the updated range maps.

### Section 3: Priors for the bMSOM for the West Andes avifauna

Vague priors cause problems in occupancy models for modest-sized datasets as they induce pushforward densities on the probability scale that concentrate near probabilities of zero and one (Lele 2015, Northrup & Gerber 2018). We specify principled priors elicited as described below. To ensure that covariate relationships are due to the data and not to our prior model we use zero-centered normal distributions for all coefficients, and we chose weakly informative prior scale parameters to understate the certainty of our prior knowledge while simultaneously avoiding outlandishly concentrated pushforward densities near zero and especially near one (occupancy probabilities may well approach zero, as in the case of strict forest specialists at pasture points, but should rarely approach one). Prior elicitation relied on JBS’s experience conducting bird surveys in Peru (Socolar et al 2019), previous trait-based modeling of neotropical bird declines (Socolar & Wilcove 2019), and previous spatially explicit mapping of neotropical bird territories (Terborgh et al 1990).

To formulate consistently interpretable priors, we standardized most covariates. However, for species-standardized elevation we divided by the standard deviation without subtracting the mean, and for the geographic range covariate we performed no additional transformation beyond what is described in the main text. This ensures that the intercept in our model corresponds to occupancy in the core of a species range. To ensure that pushforward densities are not sensitive to the arbitrary choice of reference category for dummy-coded binary predictors, we code all binary predictors as −1/1 rather than 0/1 (Agresti 2018).

#### Occupancy: intercept

We entertained a non-zero-centered prior for the intercept because zero-centered priors yielded unreasonable pushforward densities on the occupancy probability. Two independent lines of evidence imply that the intercept is substantially less than zero in our model. First, the average occupancy probability, even in the elevational and geographic core of a species’ range, is expected to be substantially less than 0.5. Less than half of the regional lowland avifauna occurs on a 97-hectare bird census plot in Peru, the best-characterized neotropical forest bird community, and within the plot only about half of species occur at any given point (Terborgh et al 1990, Walker et al 2006). This suggests an average occupancy probability of roughly 0.25. Second, we expect a substantial excess of probabilities near zero (even in the core of species’ ranges), corresponding to species-point pairs that are mismatched for habitat. However, we expect a much smaller excess of probabilities near one (spatial effects might induce some excess of probabilities near one for species with very high spatial autocorrelations, but this excess should be much smaller than the excess of near-zero probabilities). Such asymmetry is achievable in the prior pushforward density only via an intercept that is not zero-centered.

If the average occupancy probability is roughly 0.25 in our sample, then by Jensen’s inequality the average logit-probability is less than logit(0.25) = −1. If the variance in the logit-occupancy is large (due to covariate relationships and/or random effects), then the average logit-occupancy will be much less than −1. We were *a priori* certain that the variance in logit-occupancy probabilities is large, but highly uncertain about its magnitude. After defining our priors on the remaining covariates (see below), we examined how our prior choice for the intercept impacts the pushforward density for the occupancy probability of a “typical” species with covariate values near one standard deviation for every predictor. We initially experimented with a prior of Normal(−3,1) as a prior that 1) allows low values for the intercept; 2) allows values up to about −1, which is an upper bound on the plausible intercept values; and 3) results in an acceptable (rather than outrageously large) excess of probabilities near one.

However, model fitting revealed prior-data conflict, with estimates in the range of −7 to −5. Therefore, we implemented a prior of Normal(−7, 2.5) as a prior that 1) allows for extremely low values for the intercept; and 2) allows values up to about −1 without placing excess tail probability on values much greater than −1, thereby remaining consistent with the domain expertise that we had elicited previously.

#### Occupancy: range effects

In our model, two covariates (relative-elevation-squared and distance-to-range) determine whether a species is in-range or out-of-range, and we expect large effect sizes for the associated coefficients. In the context of a large effect size for relative-elevation-squared, we also entertain large effect sizes for the linear relative-elevation term or the interactions of the linear and squared relative-elevation terms with lowland occurrence. We expect these parameters to be strongly identified in the model (and confirm *a posteriori* that they are indeed strongly identified), mitigating the problems associated with weak priors. Furthermore, these coefficients have no influence on the pushforward distributions for species in the core of their geographic and elevational range. Therefore, we use very weak Normal(0,5) priors for these five coefficients.

#### Occupancy: pasture effect

We expected that the change in point-scale species richness between forest and pasture should be on average smaller than 10-fold. In the limit of low occupancy probability, this ratio corresponds to a change in logit-occupancy of about 2.3, but the corresponding effect size is larger under progressively higher baseline occupancy probabilities. Under the very high baseline occupancy probability of 0.8, this ratio corresponds to a change in logit-occupancy of roughly 4, corresponding to an effect size of 2 under our effects-coding of the interaction. We use a prior of Normal(0,1).

#### Occupancy: trait effects

Species traits might influence overall occupancy in our dataset if they are predictive of restriction to undersampled habitats, but any such effects are almost certainly not larger than the influence of traits on abundance differences between forest and pasture (see below). Aside from these effects, evidence from Peru suggests that traits are only weakly related to occupancy (Terborgh et al 1990, Russo et al 2003). Therefore, we used Normal(0, 1) priors for the main effects of traits involving habitat associations and biogeographic patterns, and Normal(0, 0.5) priors on coefficients involving body-mass or diet (see below).

#### Occupancy: trait-by-pasture interactions

Previous work in Peru suggests that the most important species traits, such as forest specialization, alter the log abundance ratio between forest and pasture by no more than about 4 (Socolar & Wilcove 2019). In the limit of low occupancy probability, this log-ratio corresponds to a change in logit-occupancy of 4, but the corresponding effect size is larger under progressively higher baseline occupancy probabilities. Under the very high baseline occupancy probability of 0.8 in preferred habitat, this log-ratio corresponds to a change in logit-occupancy of roughly 7, corresponding to an effect size of 1.75 under our effects-coding of the interaction.

Other traits have universally smaller effects, with traits related to habitat and biogeography consistently showing larger effects than traits related to body mass or diet. Therefore, we used Normal(0,1) priors for traits involving habitat or biogeography, and Normal(0,0.5) priors for traits involving body mass or diet.

#### Occupancy: random effect standard deviations

For terms with taxonomic effects of species and family, we found it easier to reason about the magnitude of the total taxonomic variation (i.e. the combined species- and family-level variation) as opposed to the species- and family-level variation in isolation. Likewise, for terms with spatial effects of both cluster and subregion, we found it easier to reason about the magnitude of the total spatial variation (i.e. the combined cluster- and subregion-level variation). Therefore, we specified these random effects by placing a half-normal prior on the square root of the combined variance, and then additively partitioning the combined variance into the two levels using a multiplicative parameter p (and its complement, one minus p), on which we placed a Uniform(0,1) prior.

To choose a prior for the standard deviation of the combined species- and family-level random intercepts, we examined our expectations for the range of typical occupancy probabilities for different species in preferred habitat (forest/pasture) in the core of their range. We felt that an appropriate distribution should be capable of covering probabilities whose logits span a range of at least 9 (e.g. probabilities as low as 1e-3 for species like *Geotrygon violacea* and *Myrmornis torquata* and as high as 0.9 for species like *Tangara chilensis* and *Zonotrichia capensis*). Thus, we chose a standard deviation of 2 for the half-normal prior on the square root of the taxonomic variance.

To choose a prior for the standard deviation of the combined species- and family-level random slopes for pasture, we considered the residual variation in responses to pasture that we would expect to find within a cohort of species with similar trait values. For example, the cohort of species that are classified as forest-present but not forest-specialist includes *Tyrannus melancholicus*, which we expected would be abundant in pasture (occupancy probabilities in excess of 0.5) and nearly absent from forests (occupancy probabilities well below 0.01), and also species like *Coeligena helianthea*, which we expected would be substantially commoner in forest than in pasture. Thus, we again felt that the random effect distribution should be capable of covering probabilities whose logits span a range of about 9, which is comfortably covered by a half-normal prior with standard deviation 1 (due to the effects coding of forest vs. pasture).

Our prior expectations were weakest for the standard deviation of the combined cluster- and subregion-level random intercepts. We felt that species modeled as present with moderate probability might be locally absent with very high probability due to the influence of unmeasured habitat features (e.g. edaphic variables). Likewise, we felt that species modeled as absent with very high probability might occur in a highly spatially correlated fashion, with high probabilities of point-level occupancy conditional on occupancy of a cluster. Therefore, we selected a very weak half-normal(0, 3) prior on the square root of the spatial variance.

We chose weak Normal(0,2) priors for the species-level slopes for relative-elevation and squared-relative-elevation. We chose these weak priors in the context of our expectation for large effect sizes in these terms and the fact that these terms do not influence the pushforward density for occupancy probabilities in the core of species ranges (see *Occupancy parameters: range effects* above).

#### Detection: intercept

Based on direct experience surveying neotropical birds on spot-mapped plots where the territories were known, we *a priori* expected average detection probabilities to be much less than 0.5, since even at the height of the dawn chorus a ten-minute point-count always detects less than half of the species present. At the same time, we felt that average values below 0.05 are *a priori* implausible (but note that due to Jensen’s inequality this lower bound estimate might correspond to an average logit substantially less than logit(0.05) = −2.9). By examining the influence of the intercept on the average of the pushforward density for the detection probability based on the full prior uncertainty for the remaining detection parameters, we found that an average probability of 0.05 is obtained when the intercept is near −6, but this estimate is expected to be substantially too low because our priors intentionally overstate the prior uncertainty in the remaining parameters. Therefore, we use a prior of Normal(−3,1) on the intercept.

#### Detection: pasture effect

We expected the effect of pasture to combine two countervailing effects. On one hand, sight lines are longer and sound propagates better in pasture than in forest. On the other hand, birds with territories that span both forest and pasture might enter the pasture portion of their territories (e.g. to forage in isolated trees; Boesing et al 2021) only infrequently. We thought that either of these effects might be large, potentially causing average detection probabilities to swing across most of their *a priori* plausible range. Due to our effects coding, we use a prior of Normal(0, 0.75) to cover such a swing.

#### Detection: time and elevation effects

Based on direct experience surveying neotropical birds, we expected detection to decay substantially though the morning in the lowlands, but not necessarily in the highlands, with effect sizes as large as a roughly ten-fold change in probability, corresponding with logit-scale changes on the order of 3 over covariate values of +/− 2-sigma. To understate our prior certainty, we used priors of Normal(0, 0.5) for the effects of time, median elevation, and the interaction between the two.

#### Detection: observer effects

Because all observers recorded sound continuously during point-counts and consulted *inter se* to identify unknown sounds, we expected observer effects to be small. We used a prior of Normal(0, 0.25) for observer effects.

#### Detection: trait effects

Previous work (in a boreal system) suggests that trait-based covariates have modest influences avian on detection probabilities (Sólymos et al 2017). We used a prior of Normal(0, 0.5) for all traits except migratory status. Because overwintering migrants are expected to be potentially much less vocal than other taxa, we allow for a larger effect size of migratory behavior, with a prior of Normal(0, 1).

#### Detection: random effect standard deviations

As for occupancy, we thought about random effect variances for detection in terms of a combined “taxonomic variance” that subsumes the species- and family-level variances.

For the intercept, we thought that detection might vary across taxa over a range of approximately 0.001 (e.g. *Neomorphus, Harpia, Nothocrax*) to 0.8, or approximately 9 units of logit-probability. We chose a half-normal prior with standard deviation 2 for the square root of the taxonomic variance.

We expected the effect of pasture to combine two countervailing effects whose importance might vary by species (see *Detection: pasture effect* above). Therefore, we thought that the random-effect variation might potentially exceed our prior uncertainty in the mean effect size, and so we chose a half-normal prior with standard deviation 1 for the square root of the taxonomic variance.

For species-specific variation in time-of-day relationships, we expected substantial variation between species that regularly join the dawn chorus versus species that rarely sing but are often detected in mixed flocks that form later in the morning. Therefore, we use a half-normal prior with standard deviation 1 on the standard deviation of the species effect variance.

### Section 4: The saga of the data-augmented model

We attempted to fit the data-augmented model under the same priors as the bMSOM for all shared parameters, replacing the species-standardized elevations with scaled raw elevations and placing a uniform prior on *ω*, the probability that a given species is included in the metacommunity. However, model fitting proved challenging. During the early phase of warmup all MCMC chains entered a region of the posterior where they consistently saturated the treedepth and failed to mix quickly. As warmup proceeded, the chains adapted to the geometry of this region of the posterior but failed to explore the bulk of the posterior. Every chain entered a region where the standard deviation of the species-specific random intercept for either occupancy or detection approached zero (e.g. values less than 10^−10^) and then slowly recovered. Yet this recovery was so glacially slow that it quickly became clear that we could not achieve correct inference using our available computational resources except by modifying our approach.

Fortunately, all chains agreed in estimating extremely high values of *ω*, essentially equal to one. We noted that if we fix *ω* to 1, the data-augmented MSOM becomes equivalent to the traditional MSOM fit to the augmented dataset. On the hypothesis that *ω* would remain near one at convergence in the data-augmented model, we fit the traditional MSOM to the augmented data to learn the approximate location and scale of the posterior. The traditional model ran much faster. Because our main purpose was to obtain useful estimates for the inverse metric and for initial values in the data-augmented model, we ran four chains until one chain completed its 1000 warmup iterations, at which point we terminated the computation.

We then extracted the inverse metric from the chain that finished warmup as well as the final iteration from all four chains, and we again attempted to fit the data-augmented MSOM, this time initializing the inverse metric using the diagonal metric extracted from the traditional MSOM and initializing the chains at the final iterations from each of the four chains from the traditional MSOM. We initialized the diagonal inverse metric entry for *ω* to Stan’s default value of one, and the intercept for *ω* at 0.990. To avoid bad early updates to the inverse metric, we also specified that the initial adaptation window for computing the inverse metric should run for 50 iterations rather than the default 25. To avoid unnecessarily long integration times after warmup we specified that the term buffer should run for 100 iterations rather than the default 50. This second attempt at fitting data-augmented MSOM immediately showed much better mixing behavior than our initial attempt, but still took roughly a week per chain to run. One of the four chains was slightly less successful in its adaptation and adapted to a step-size that required an increase in treedepth from 7 to 8 and therefore twice the computation time. We terminated this chain. At the end of model fitting, r-hat values for some parameters (computed over the three chains that were retained) were problematic (as high as 1.16). However, all three chains again agreed that *ω* was very high (95% credible interval 0.9977–1.0000, r-hat = 1.01). The chains also agreed that the occupancy intercept was extremely low (95% credible interval −13.8 – −16.8, r-hat = 1.04), which is low enough to substantially conflict with our already-low prior of Normal(−7.5, 2.5).

Thus, in a final step, we switched back to the traditional MSOM, confident that the data-augmented model truly yielded *ω* values near unity, and we replaced the Normal(−7.5, 2.5) prior on the occupancy intercept with a logistic prior (i.e. a flat prior on the probability scale), which decays marginally slower than Normal(−7.5, 2.5) in the region of −13.8 – −16.8. To avoid spending excessive computational resources on warmup, we terminated this model after one chain finished warmup and then ran four new chains with no warmup using initial values taken from the final state of the previous chains, and inverse metric and step size taken from the chain that finished warmup. Even still, parts of this model did not converge, especially the detection intercept (rhat = 1.14) and species-specific parameters related to detection (rhat as high as 1.16). The lack of convergence likely reflects the inherent misspecification of the data-augmented model for this system. Nevertheless, all hyperparameters in the occupancy sub-model were somewhat better behaved (maximum r-hat = 1.04) and so we used this final model as the basis for tentative inference on patterns of species richness.

## Supplementary Tables

**Table S1.**
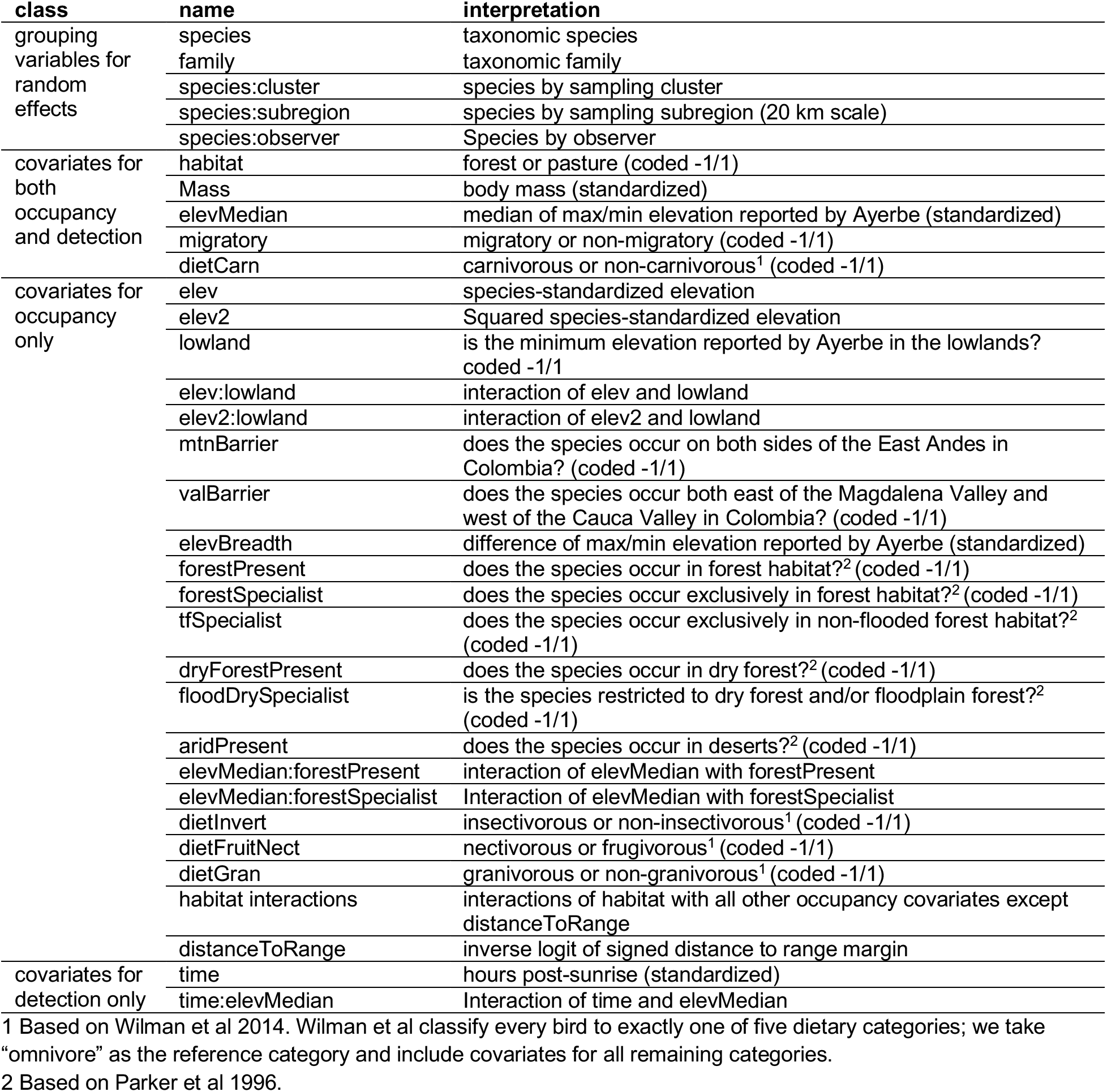
Model covariates and grouping terms for the West Andes

**Table S2.** Parameter estimates for hyperparameters and fixed effects in the West Andes. (Continued on next page).

| class | name | grouped by | mean | lower CI<sup>1</sup> | upper CI<sup>1</sup> |
| --- | --- | --- | --- | --- | --- |
| fixed effects<br>on occupancy | intercept |  | -7.27 | -9.59 | -4.99 |
|  | elev |  | -3.67 | -5.10 | -2.30 |
|  | elev2 |  | -5.44 | -6.91 | -4.09 |
|  | habitat |  | 1.49 | 0.26 | 2.75 |
|  | distanceToRange |  | -3.50 | -4.83 | -2.24 |
|  | lowland |  | -0.22 | -0.92 | 0.50 |
|  | elev:lowland |  | -4.63 | -6.01 | -3.26 |
|  | elev2:lowland |  | 0.70 | -0.52 | 1.81 |
|  | mtnBarrier |  | 0.28 | -0.35 | 0.91 |
|  | valBarrier |  | -0.21 | -0.96 | 0.55 |
|  | elevMedian |  | -0.70 | -1.71 | 0.29 |
|  | elevBreadth |  | -0.03 | -0.69 | 0.63 |
|  | forestPresent |  | 1.74 | 0.84 | 2.65 |
|  | forestSpecialist |  | 0.81 | 0.16 | 1.46 |
|  | tfSpecialist |  | 0.37 | -0.12 | 0.87 |
|  | dryForestPresent |  | -0.07 | -0.64 | 0.51 |
|  | floodDrySpecialist |  | -0.09 | -1.39 | 1.18 |
|  | aridPresent |  | -0.40 | -1.43 | 0.61 |
|  | migratory |  | -0.93 | -2.19 | 0.39 |
|  | elevMedian:forestPresent |  | 0.60 | -0.37 | 1.58 |
|  | elevMedian:forestSpecialist |  | 0.38 | -0.26 | 1.03 |
|  | mass |  | -0.74 | -1.34 | -0.14 |
|  | dietInvert |  | -0.33 | -0.80 | 0.13 |
|  | dietCarn |  | -0.57 | -1.35 | 0.21 |
|  | dietFruitNect |  | 0.34 | -0.13 | 0.82 |
|  | dietGran |  | -0.06 | -0.67 | 0.55 |
|  | habitat:mtnBarrier |  | -0.24 | -0.65 | 0.15 |
|  | habitat:valBarrier |  | -0.32 | -0.79 | 0.16 |
|  | habitat:elevMedian |  | -0.64 | -1.47 | 0.16 |
|  | habitat:elevBreadth |  | 0.71 | 0.32 | 1.10 |
|  | habitat:forestPresent |  | -0.45 | -1.18 | 0.23 |
|  | habitat:forestSpecialist |  | -2.03 | -2.48 | -1.59 |
|  | habitat:tfSpecialist |  | 0.21 | -0.08 | 0.51 |
|  | habitat:dryForestPresent |  | 0.60 | 0.23 | 0.97 |
|  | habitat:floodDrySpecialist |  | 0.03 | -0.91 | 1.00 |
|  | habitat:aridPresent |  | 0.51 | -0.33 | 1.39 |
|  | habitat:migratory |  | 0.37 | -0.39 | 1.12 |
|  | habitat:elevMedian:forestPresent |  | 0.32 | -0.52 | 1.16 |
|  | habitat:elevMedian:forestSpecialist |  | 0.32 | -0.08 | 0.72 |
|  | habitat:mass |  | -0.12 | -0.45 | 0.21 |
|  | habitat:dietInvert |  | -0.34 | -0.64 | -0.04 |
|  | habitat:dietCarn |  | -0.48 | -1.18 | 0.21 |
|  | habitat:dietFruitNect |  | -0.02 | -0.33 | 0.29 |
|  | habitat:dietGran |  | -0.24 | -0.26 | 0.74 |
|  | distanceToRange |  | -3.50 | -4.83 | -2.24 |
| occupancy<br>random effect<br>standard<br>deviations | sd_intercept_taxonomic | species, family<sup>2</sup> | 3.99 | 3.22 | 4.83 |
|  | p_intercept_taxonomic | proportion species<sup>3</sup> | 0.63 | 0.42 | 0.84 |
|  | sd_intercept_spatial | cluster, subregion<sup>2</sup> | 2.81 | 2.54 | 3.11 |
|  | p_intercept_spatial | proportion cluster<sup>3</sup> | 0.63 | 0.56 | 0.70 |
|  | sd_relev | species | 2.85 | 2.21 | 3.47 |
|  | sd_relev2 | species | 2.66 | 2.02 | 3.38 |
|  | sd_habitat_taxonomic | species, family<sup>2</sup> | 1.57 | 1.26 | 1.90 |
|  | p_habitat_taxonomic | proportion species<sup>3</sup> |  |  |  |
|  | sd_distanceToRange | species | 3.69 | 1.63 | 5.30 |
| fixed effects<br>on detection | intercept |  | -4.50 | -5.41 | -3.56 |
|  | habitat |  | -0.07 | -0.22 | 0.08 |
|  | time |  | -0.20 | -0.25 | -0.16 |
|  | mass |  | -0.41 | -0.66 | -0.16 |
|  | elevMedian |  | -0.02 | -0.24 | 0.18 |
|  | migratory |  | -1.35 | -1.87 | -0.79 |
|  | dietCarn |  | -0.62 | -1.31 | 0.12 |
|  | time:elevMedian |  | -0.12 | -0.17 | -0.06 |
| detection<br>random effect<br>standard<br>deviations | sd_intercept_taxonomic | species, family<sup>2</sup> | 1.21 | 1.02 | 1.42 |
|  | p_intercept_taxonomic | proportion species<sup>3</sup> | 0.85 | 0.66 | 0.98 |
|  | sd_intercept_spObs | species:observer | 0.23 | 0.16 | 0.30 |
|  | sd_habitat_taxonomic | species, family<sup>2</sup> | 0.70 | 0.59 | 0.83 |
|  | p_habitat_taxonomic | proportion species<sup>3</sup> | 0.91 | 0.80 | 0.99 |
|  | sd_time | species | 0.20 | 0.15 | 0.24 |

1 Lower and upper bounds of a 95% credible interval (i.e. 0.025 and the 0.975 quantiles of the marginal posterior).
2 This is the square root of the combined variance associated with random effects of species and family (or cluster and subregion).
3 This is the proportion of the combined variance that is associated with species rather than family (or cluster rather than subregion).

## Supplementary Figures

**Figure S1.**
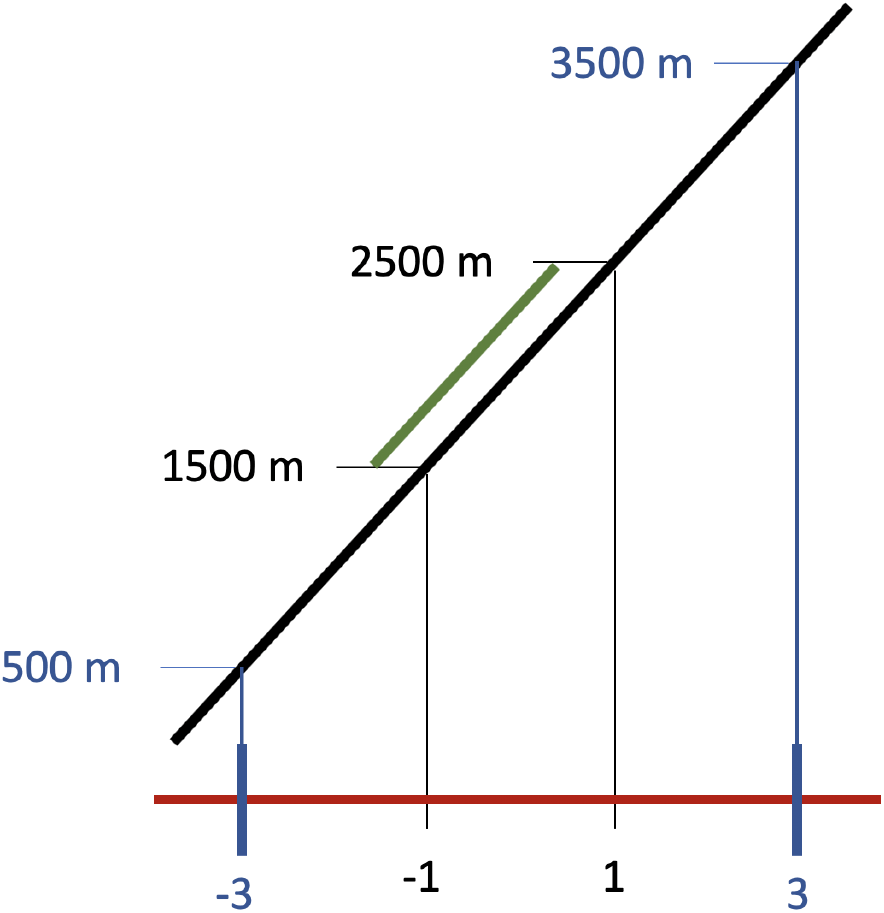
We compute species-standardized elevations for each sampling point and perform biogeographic clipping as follows. We linearly rescale the actual elevation gradient (black line) to a new species-specific gradient (red line) such that the published species-specific minimum and maximum elevations (in this example 1500 m and 2500 m) correspond to −1 and 1. We then perform biogeographic clipping at species-standardized elevations of −3 and 3, in this case corresponding to 500 m and 3500 m.

**Figure S2.**
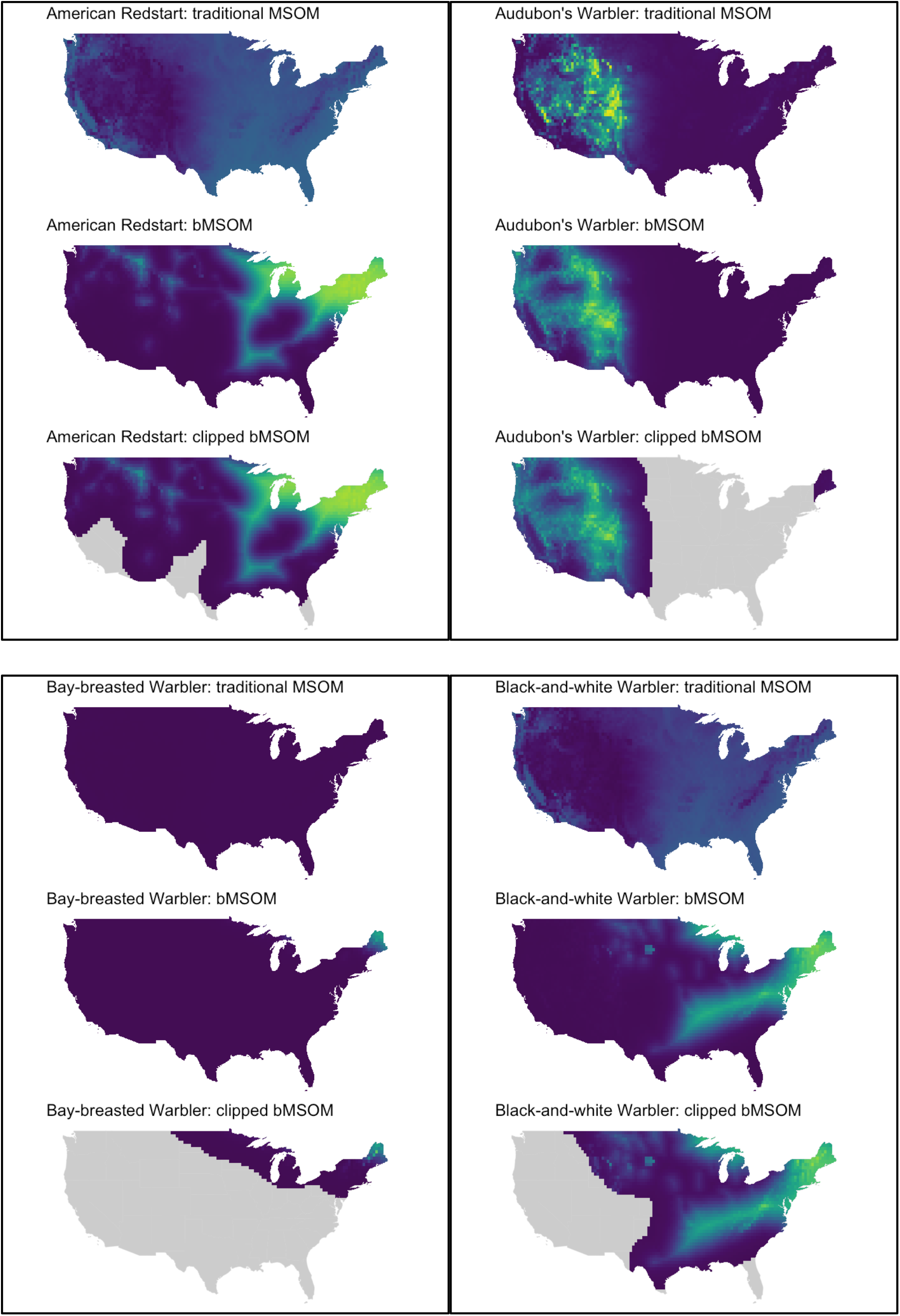
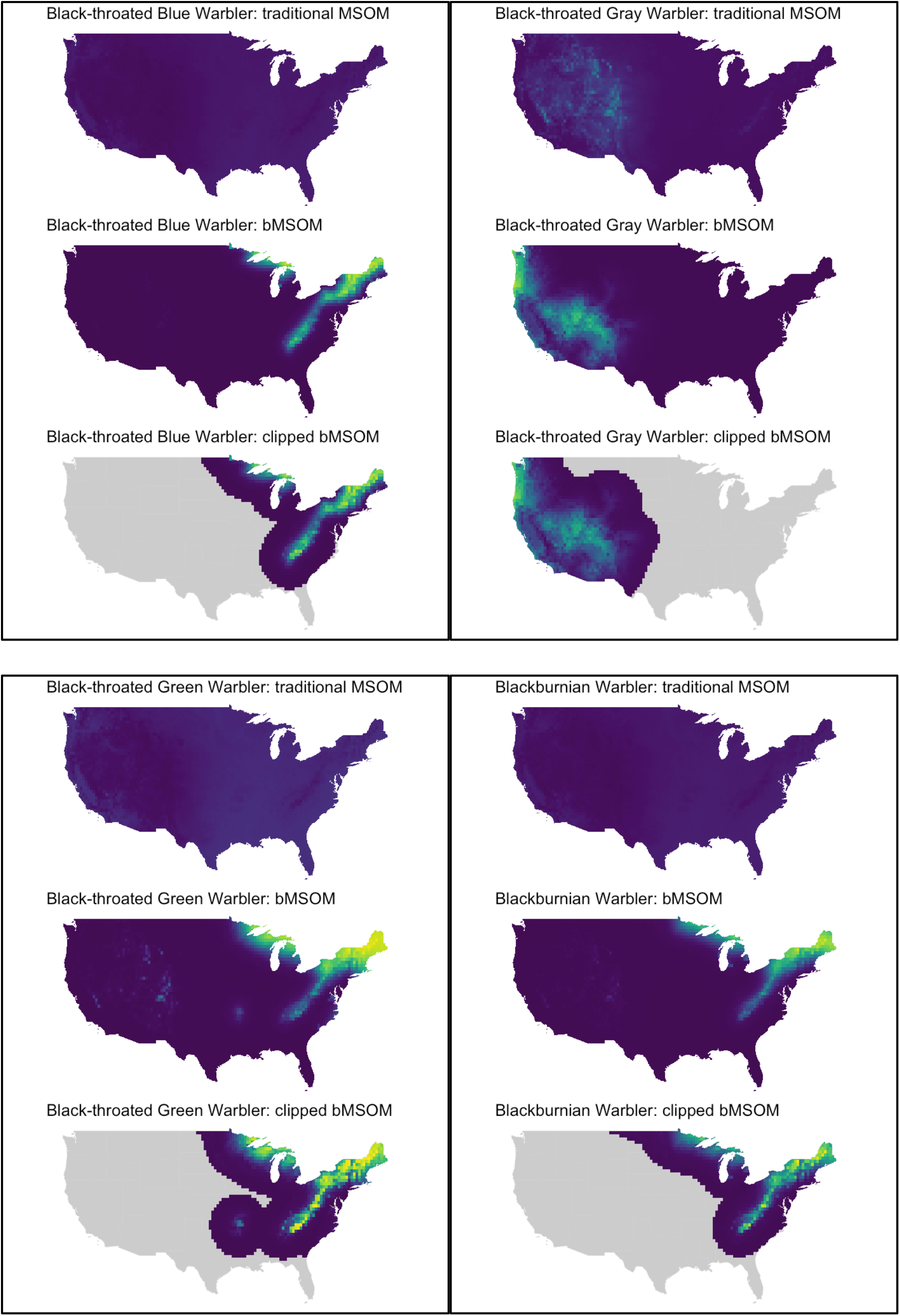
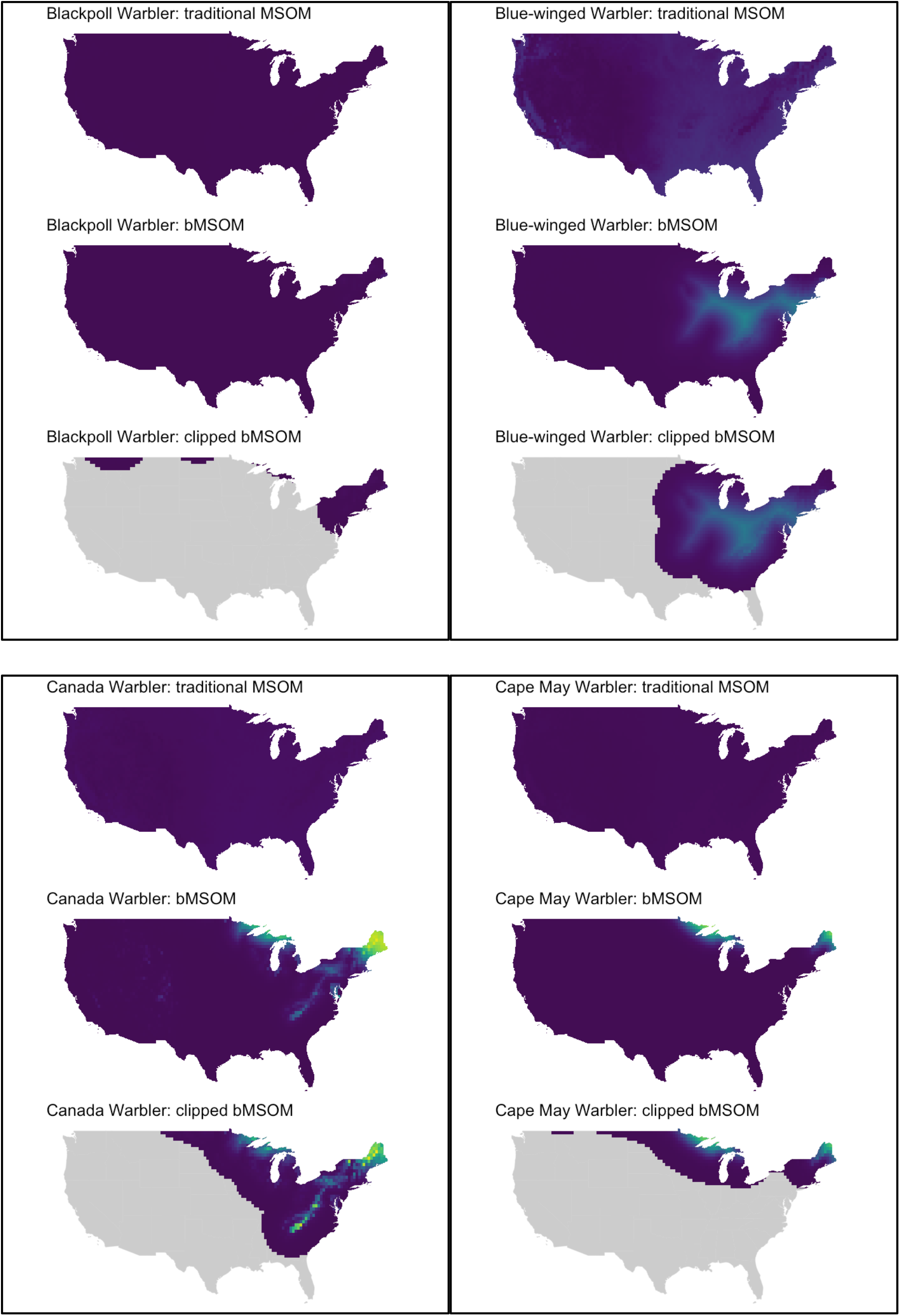
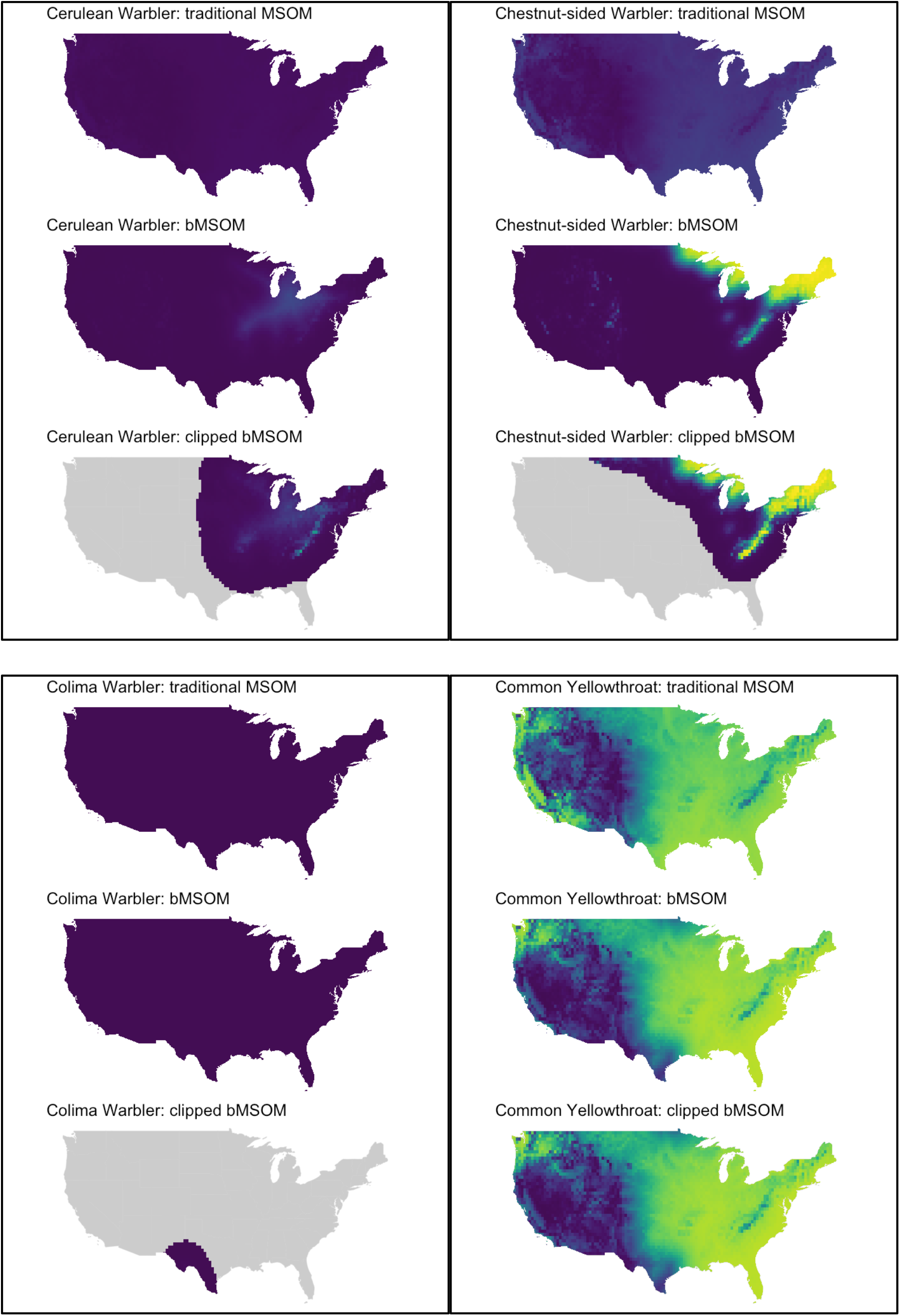
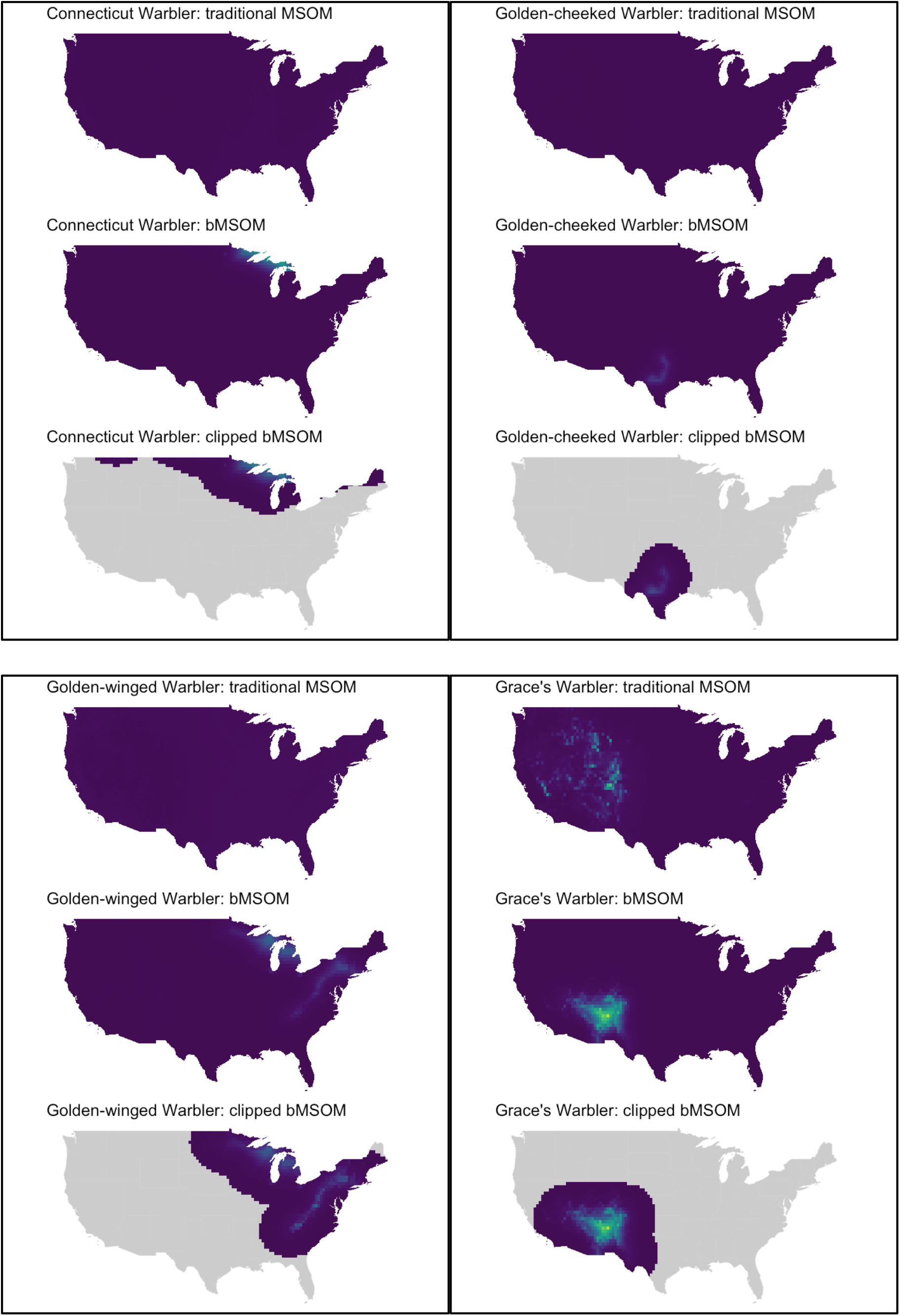
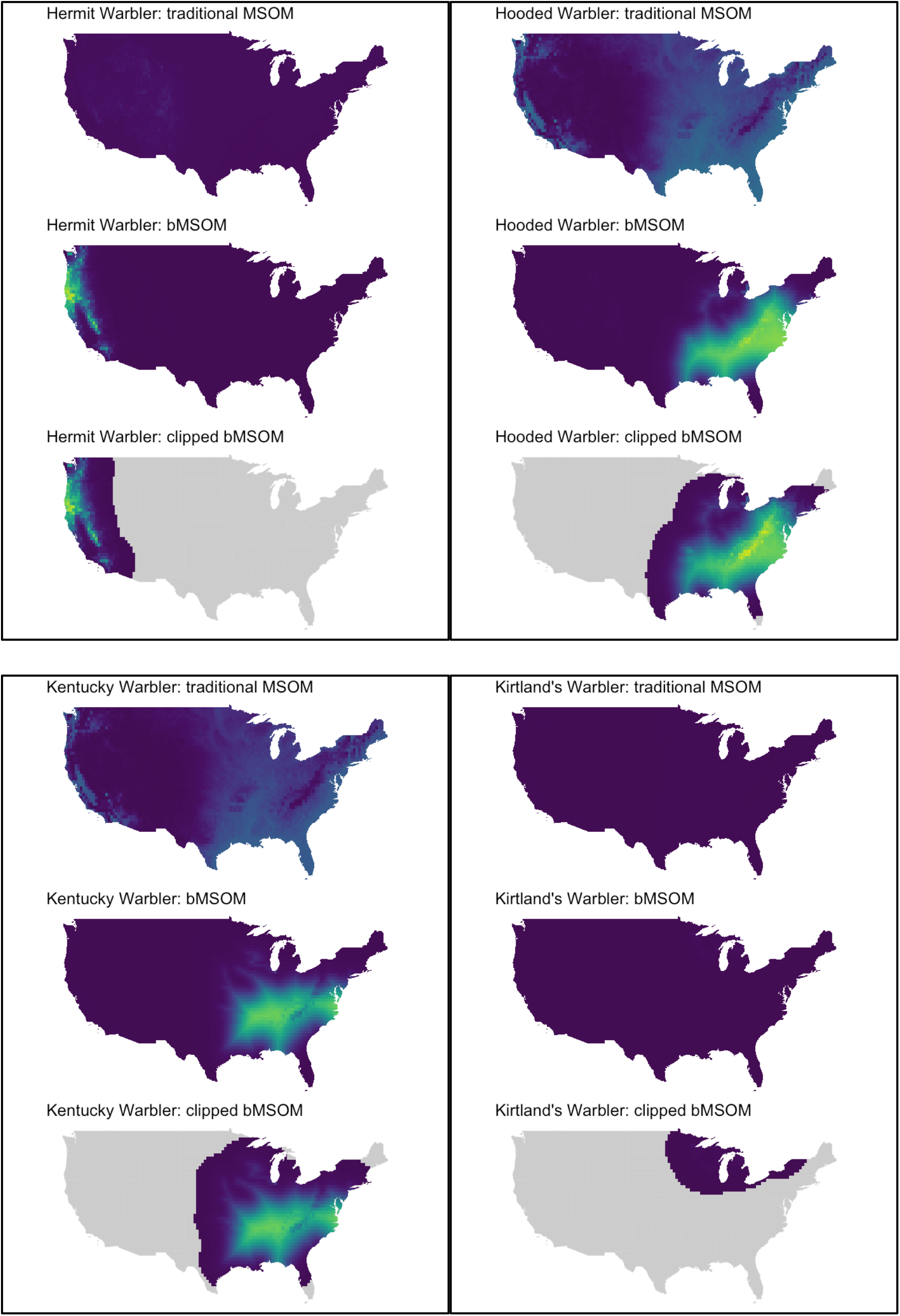
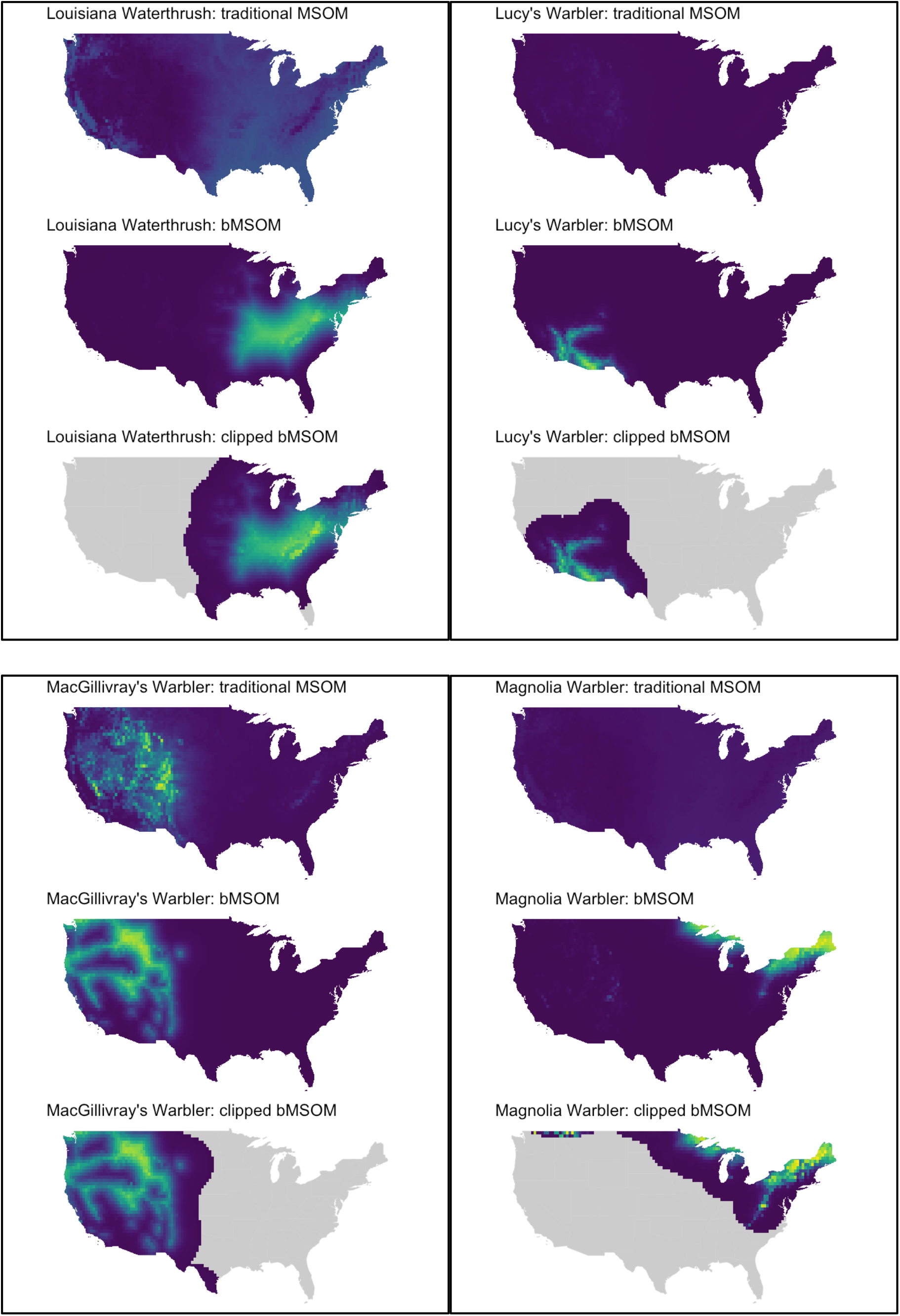
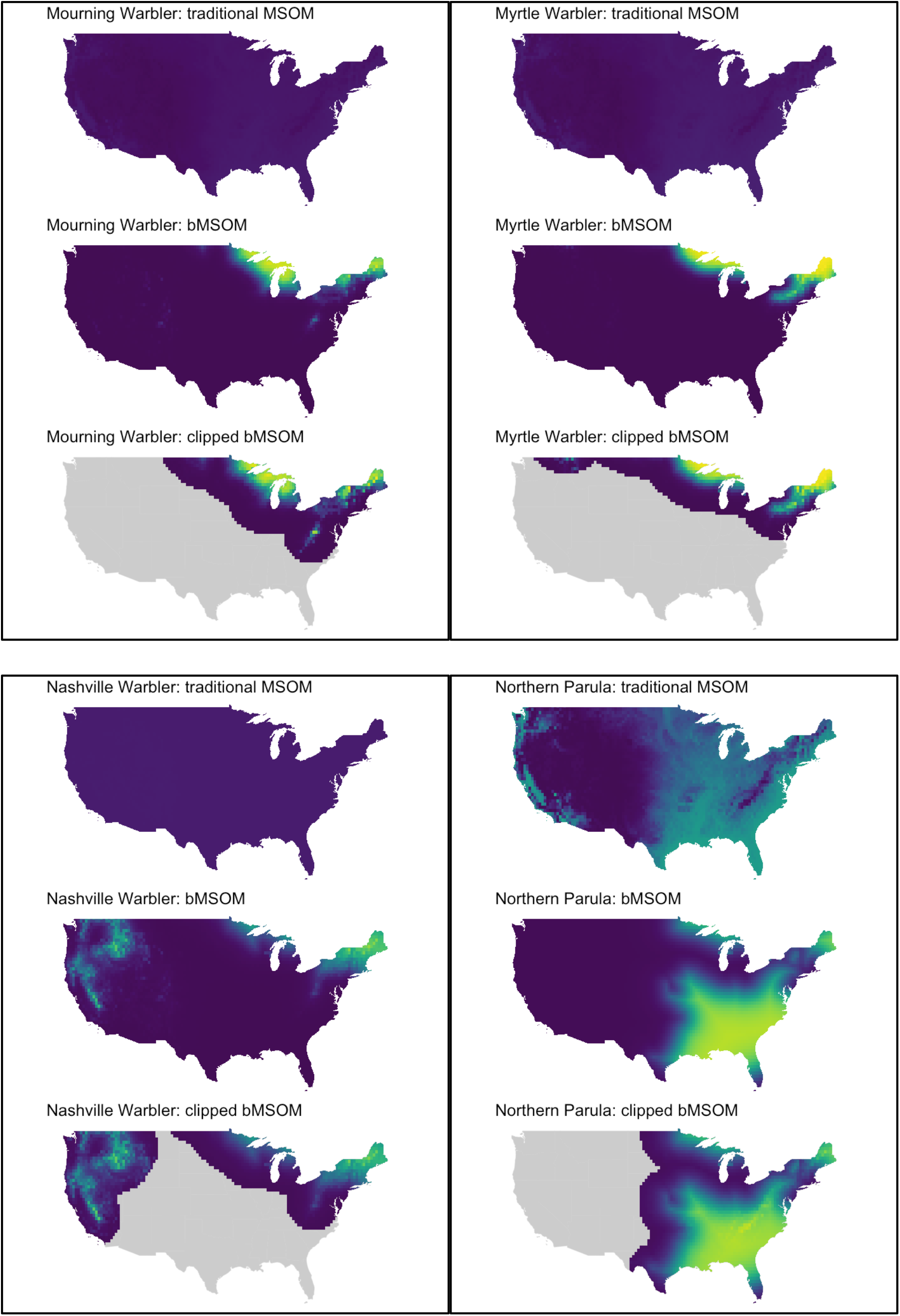
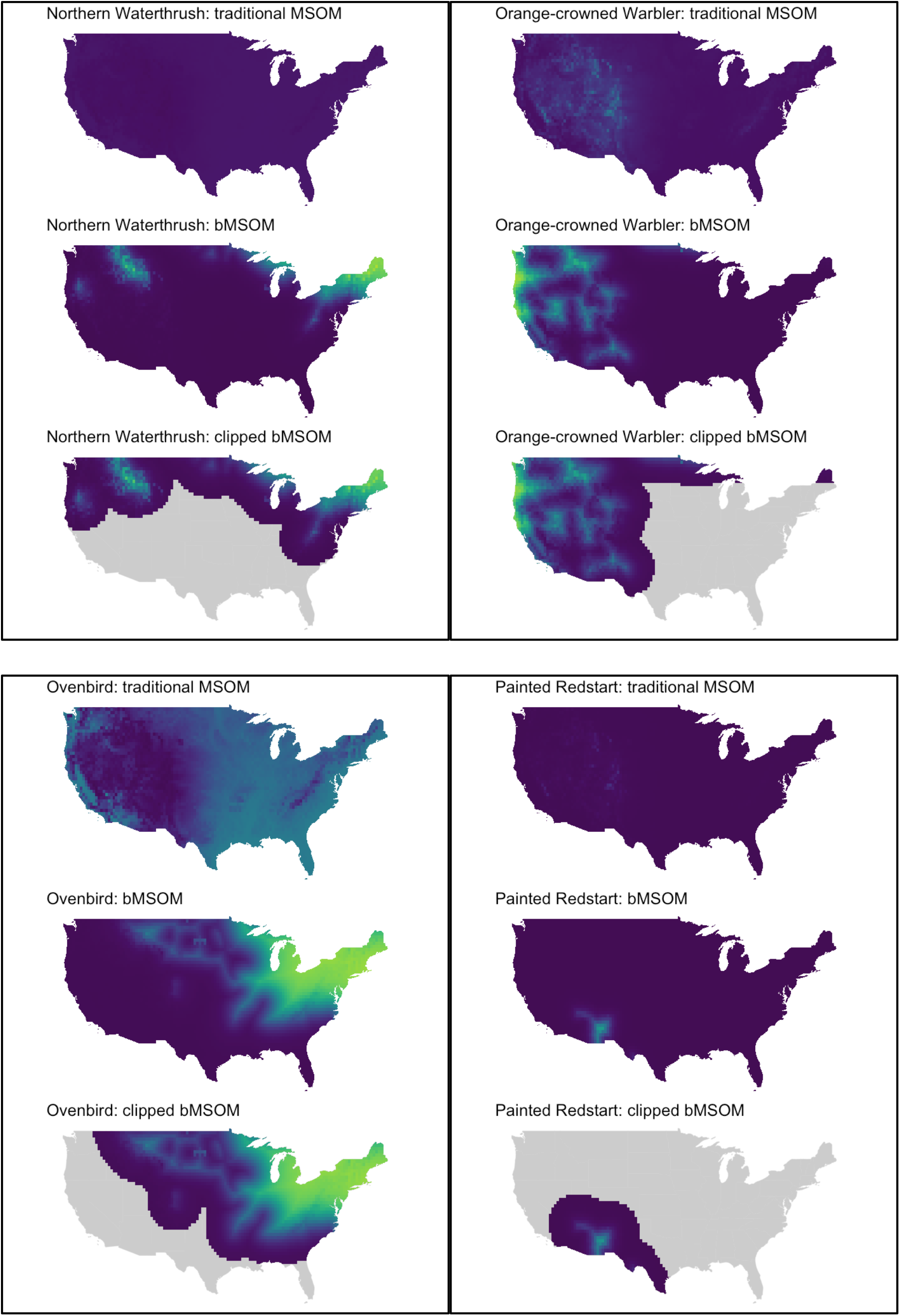
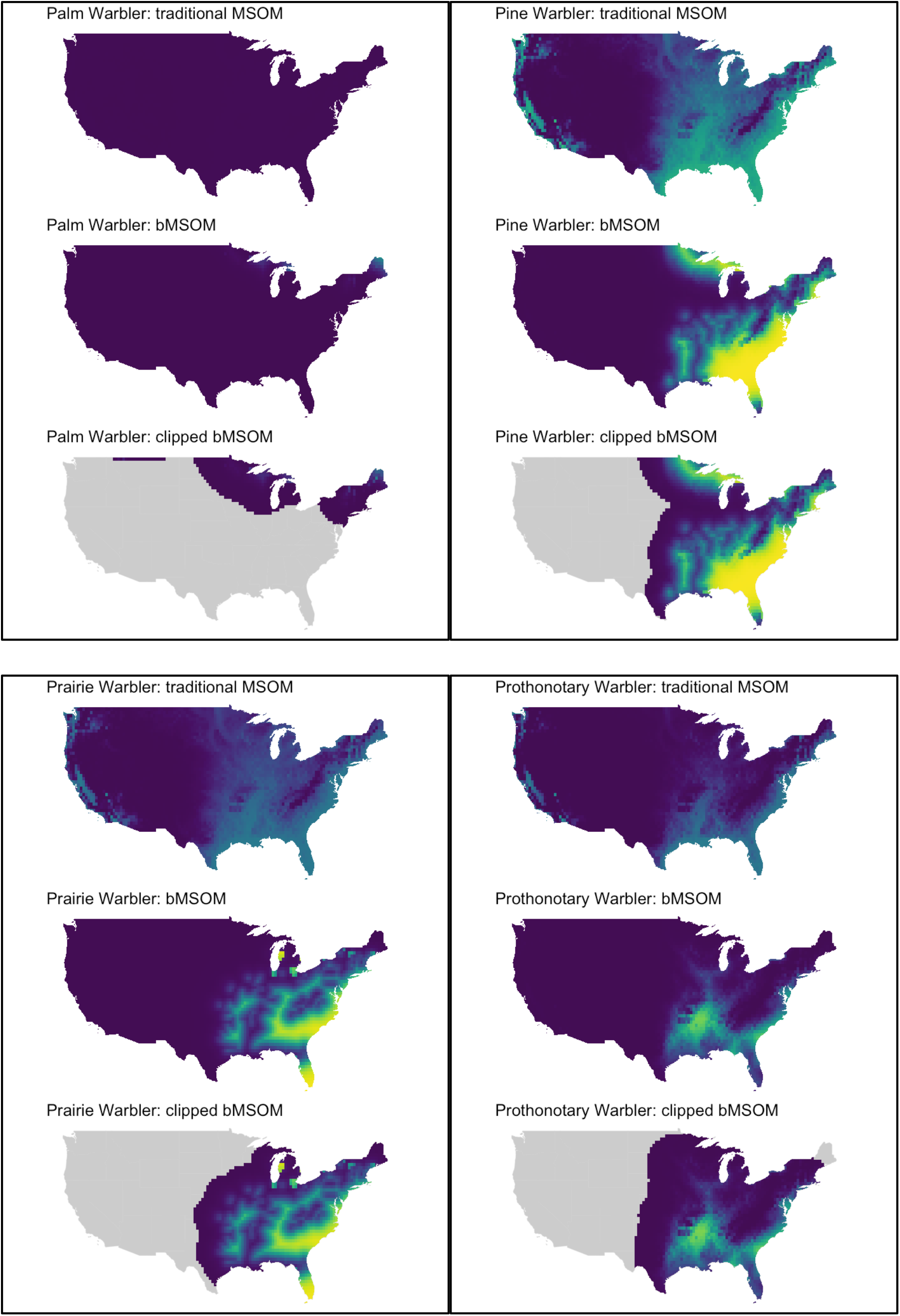
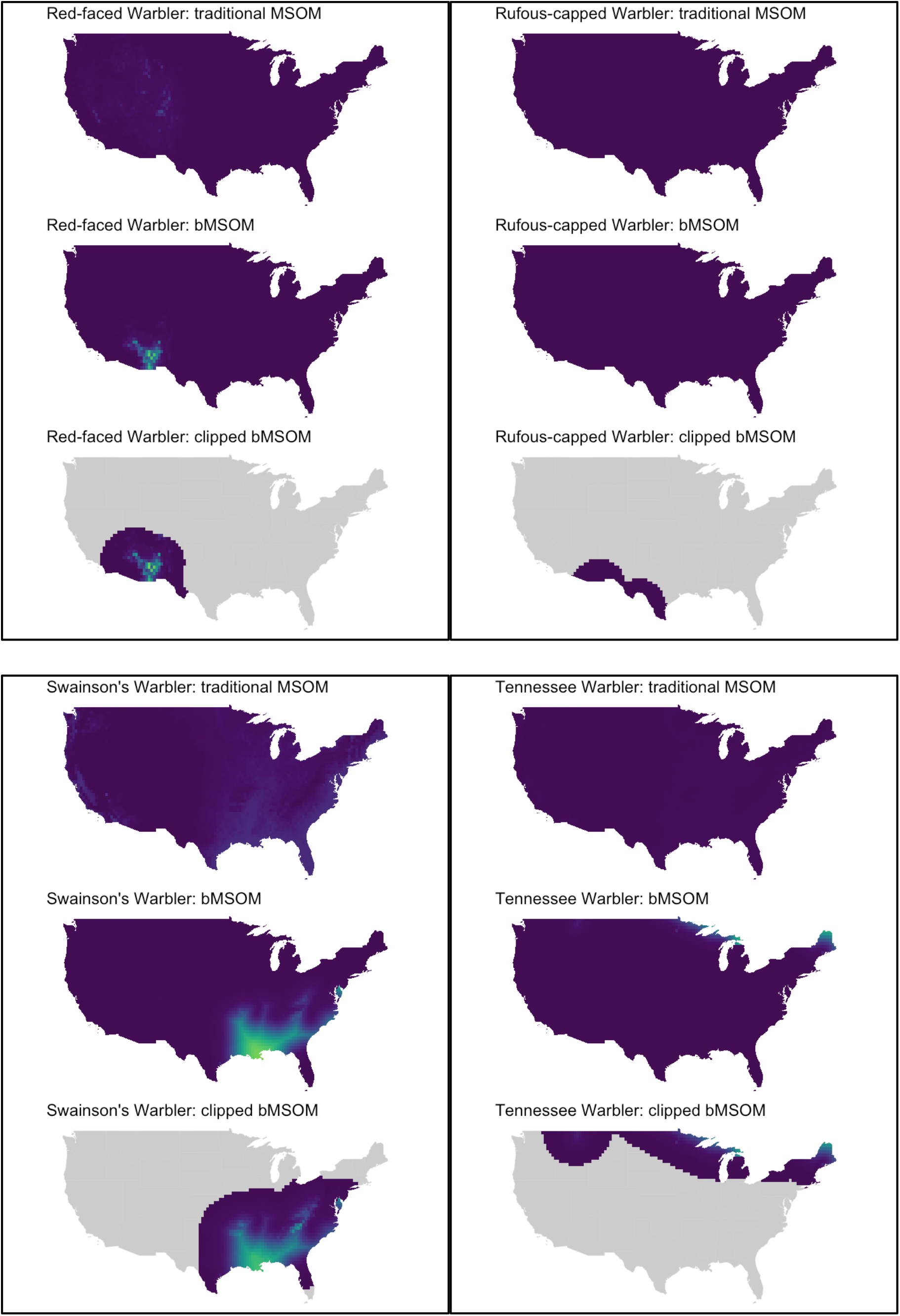
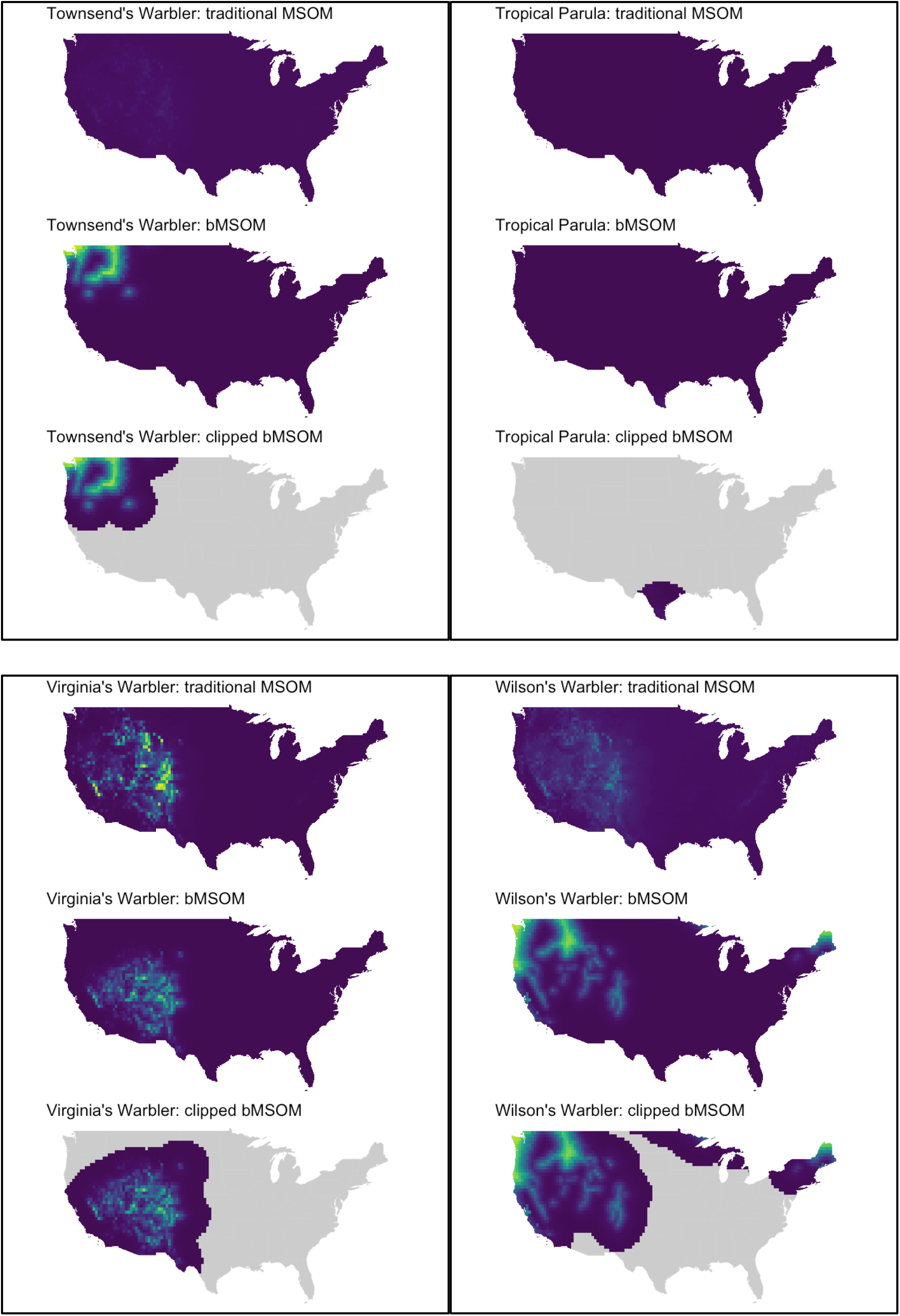
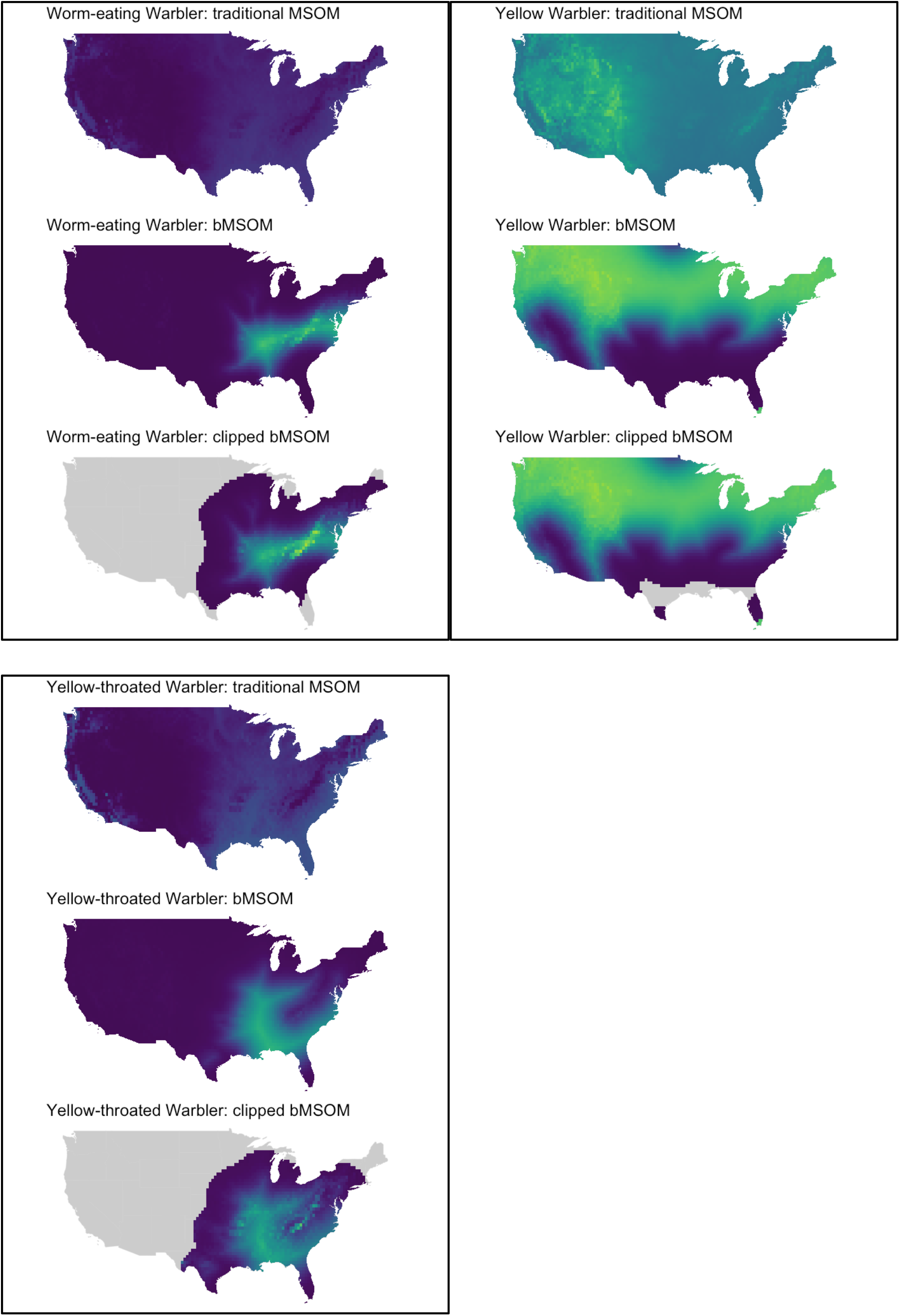
(begins on next page) Predicted occupancy probabilities across the coterminous United States for all 51 modeled warbler species based on the traditional multi-species occupancy model (traditional MSOM), the biogeographic model (bMSOM), and the clipped biogeographic model (clipped bMSOM). The color-scale is equivalent to the scale in figures 2 and 3 of the main text.

